# Tunicate metatranscriptomes reveal ancient virus-host co-divergence and inter-order recombination in the evolutionary history of disease-causing viruses

**DOI:** 10.1101/2024.12.15.628590

**Authors:** Mary E. Petrone, Joe Grove, Rhys H. Parry, Kate Van Brussel, Jonathon C.O. Mifsud, Zuhairah Dindar, Shi-qiang Mei, Mang Shi, Olivia M. H. Turnbull, Ezequiel M. Marzinelli, Edward C. Holmes

**Affiliations:** School of Medical Sciences, The University of Sydney, Sydney, NSW 2006, Australia; Laboratory of Data Discovery for Health Limited, Hong Kong SAR, China; MRC-University of Glasgow Centre for Virus Research, Glasgow G61 1QH, UK; School of Chemistry and Molecular Biosciences, The University of Queensland, Brisbane, QLD 4067, Australia; Centre for Marine Science and Innovation, Evolution & Ecology Research Centre, School of Biological, Earth and Environmental Sciences, UNSW, Sydney, NSW 2033, Australia; State Key Laboratory for Biocontrol, School of Medicine, Shenzhen Campus of Sun Yat-sen University, Sun Yat-sen University, Shenzhen, China; National Key Laboratory of Intelligent Tracking and Forecasting for Infectious Diseases, Sun Yat-sen University, Shenzhen, China; Shenzhen Key Laboratory for Systems Medicine in Inflammatory Diseases, Shenzhen Campus of Sun Yat-sen University, Sun Yat-sen University, Shenzhen, China; Guangdong Provincial Center for Disease Control and Prevention, Guangzhou, China; School of Life and Environmental Sciences, The University of Sydney, Sydney, New South Wales, Australia

**Author notes:** Correspondence: Prof. Edward C. Holmes University of Sydney.

**Keywords:** RNA virus, evolutionary virology, influenza virus, recombination, protein structure prediction

## Abstract

Tunicates are a key transitional taxon in animal evolution as the closest extant invertebrate relatives of the vertebrates. Their viruses may also reflect this transitional state. Yet, it is not known whether tunicate viruses are more closely related to vertebrate- or invertebrate-infecting viral lineages. We analysed primary and publicly available RNA libraries to extend the known diversity of tunicate-associated viruses and determine their relationship to viruses of other animals. We present evidence that influenza viruses, alphaviruses, and some mononegaviruses emerged prior to the evolution of vertebrates. We also show that the recombination of glycoproteins between different orders of RNA viruses, including between positive- and negative-sense viruses, may have shaped the evolution of multiple lineages. Our study reveals that some disease-causing RNA virus lineages were present in early chordates and highlights that the evolution of structural genes may be incongruent with that of the highly conserved RNA-dependent RNA polymerase.

## INTRODUCTION

Tunicates, sometimes referred to as urochordates, are filter-feeding marine invertebrate members of the phylum Chordata. The Tunicata subphylum comprises three classes – the *Ascidiacea*, *Thaliacea*, and *Appendicularia* – that exhibit distinct life cycles and body plans. The *Ascidiacea* (“sea squirts”) are the canonical tunicate, as it was their thick, tunic-like encasement made of cellulose that inspired the name “Tunicata” (Lamarck, 1816). Ascidians begin life as free-swimming ‘tadpole’ larvae with a simple chordate body-plan, before attaching to manmade or natural surfaces where they undergo profound morphological changes and become sessile adults. Despite a simple outward appearance, adult ascidians possess a complement of organs, including a pharynx, reproductive organs, digestive organs, neural gland, and a heart that circulates hemolymph. The *Thaliacea* (salps and doliolids) do not attach to surfaces when they mature and remain free floating at a wide range of ocean depths for the duration of their lives^1^. Like that of the ascidians, the doliolid life cycle includes a tadpole stage. In contrast to the *Ascidiacea* and *Thaliacea*, the *Appendicularia* do not undergo morphological changes as they mature, instead crafting intricate mucosal “houses” around their tadpole-like bodies to trap food^2^.

The Tunicata are of evolutionary importance because all tunicate larvae possess a notochord. It was the discovery of this feature by Alexander Kovalevsky in 1866 that prompted the reclassification of the Tunicata from Mollusca to Chordata^3^. Since then, ascidians, in particular *Ciona intestinalis* (Phlebobranchia, *Ascidiacea*), have been used to elucidate the mechanisms driving gastrulation and the formation of the notochord during the early stages of chordate development^4,5^. The precise phylogenetic relationship of the Tunicata to the Vertebrata has sparked considerable debate. While it has been argued that cephalochordates (lancelets) are the sister group of the vertebrates^6,7^, molecular data support the conclusion that tunicates are indeed the closest relatives of the vertebrates^8–10^. Thus, tunicates represent a key transitional taxon in animal evolution as the “most vertebrate-like invertebrate”.

Despite their scientific importance, very little is known about the viruses that infect tunicates aside from studies of bacteriophage diversity in tunicate gut microbiomes^11–13^ and incidental findings from metagenomic sequencing. Outbreak investigations into the cause of soft tunic syndrome, a disease that affects edible ascidians, identified a novel birnavirus^14,15^. However, a pathogenic protist was later determined to be the aetiological agent^16–18^. Data mining of tunicate metatranscriptomes identified two members of the *Medioniviridae* (*Botrylloides leachii* nidovirus, NCBI accession MK956105 and Tognidovirus botryllus22085, accession BK066634^19^) that utilise togavirus-like class II fusion proteins. Tunicate-associated Beihai orthomyxo-like virus 2 was originally classified within the *Orthomyxoviridae*^20^, but was later found to be a member of the divergent “Cnidenomoviridae” (order *Articulavirales*)^21^. Lastly, Chowder bay tunicate associated flavi-like virus (OX394159) was identified as a member of the *Flaviviridae* related to other marine invertebrate viruses among the “tamanaviruses”^22^. The diversity of tunicate-associated viruses in such a small data set hints at unrealised RNA virus diversity that has yet to be fully explored.

As well as uncovering more of the global virosphere, a thorough characterisation of the tunicate virome could address a key evolutionary question: as tunicates are the sister-group to the vertebrates, do their RNA viruses share a closer phylogenetic relationship with viruses found in vertebrate or invertebrate hosts? If the former, these vertebrate-infecting viral lineages may have predated the evolution of the Vertebrata themselves. In addition, the emergence of the vertebrates was marked by major evolutionary innovations, including the development of an adaptive immune system, which may have in turn impacted virus diversity. Hence, the identification of vertebrate-infecting-like viruses in tunicates would provide a starting point for understanding how these innovations in vertebrate evolution shaped the long-term evolutionary history of disease-causing viruses. Herein, we explored this question by systematically screening primary and publicly available tunicate metatranscriptomes for RNA viruses. We combined phylogenetic analysis and protein structural prediction to re-assess the evolutionary origins of some vertebrate-infecting viral lineages and examine instances of recombination between viral orders.

## RESULTS

### Tunicate metatranscriptomes are a rich source of viral diversity

We assembled a data set of 3,499 tunicate metatranscriptomes from publicly available SRA libraries and primary samples. The SRA libraries comprised all three classes within the Tunicata (*Ascidiacea*, *Appendicularia*, and *Thaliacea*), albeit with an uneven distribution: the majority of libraries were *Ascidiacea* (93%), and orders within each class were not uniformly represented (**Fig. 1a**). The primary samples sequenced for this study comprised 31 samples of *Ascidiacea* collected from Chowder Bay, Sydney Harbour, Australia at two time points approximately one year apart (November 2021 and October 2022) (**Fig. 1b**).

**Figure 1.**
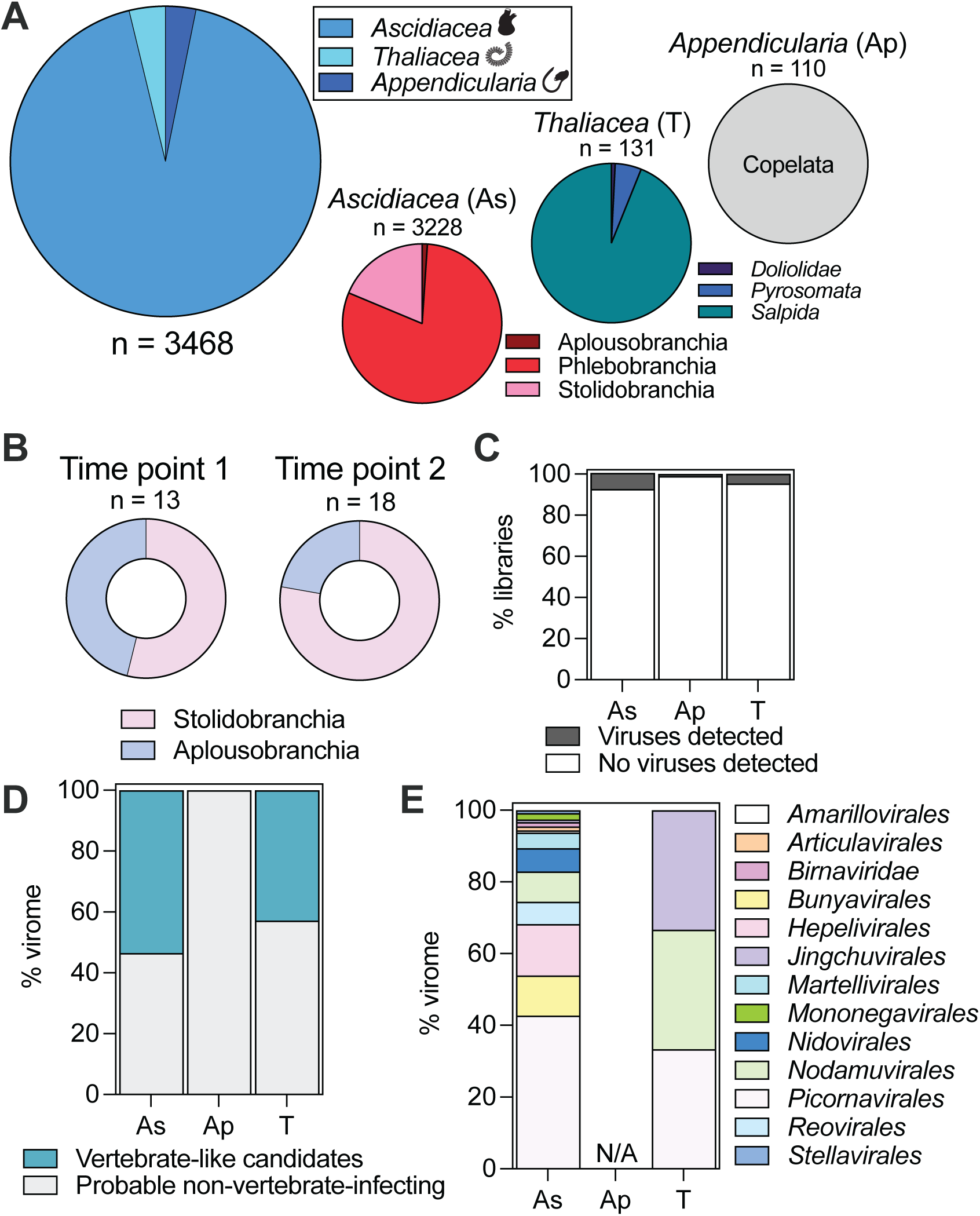
Diversity of RNA viruses identified in Tunicata metatranscriptomes. (a) Distribution of host organisms represented in SRA libraries as of February 2024. (b) Distribution of host organisms of primary samples collected in November 2021 (time point 1) and October 2022 (time point 2). (c) Percentage of libraries in which viral contigs were detected by host class. (d) Distribution of viruses belonging to orders with known vertebrate hosts (vertebrate-like candidates) and with no known vertebrate hosts (probable non-vertebrate infecting). (e) Distribution of vertebrate-like candidates in libraries by host class. As: *Ascidiacea*, Ap: *Appendicularia*, T: *Thaliacea*.

This data set yielded extensive viral diversity, particularly in the primary samples. In the SRA and primary libraries combined, we detected putative viral contigs in 242 of the 3,480 libraries analysed (7.0%) (**Fig. 1c**). However, we identified putative viral contigs in all the libraries generated from primary samples, which may be indicative of differences in viral prevalence between primary and laboratory reared organisms and library selection protocols (e.g., the use of poly-A selection in 440 libraries). Of the 1,370 putative viral contigs detected, 33 belonged to known viral taxa and an additional 187 shared sequence similarity with viruses that are unclassified by NCBI or the International Committee on Taxonomy of Viruses (ICTV). We further categorised these contigs as “vertebrate-like candidates” and “probable non-vertebrate infecting” according to the viral order to which they belonged (**Fig. 1d**). Contigs assigned to “vertebrate-like candidates” belonged to taxa with known vertebrate hosts: the *Amarillovirales, Articulavirales, Birnaviridae, Bunyavirales, Hepelivirales, Jingchuvirales, Martellivirales, Mononegavirales, Nidovirales, Nodamuvirales, Picornavirales, Reovirales,* and *Stellavirales* (**Fig. 1e**). Contigs sharing sequence similarity to viruses in taxa that have no known vertebrate hosts (e.g., the *Ourlivirales*) were classified as “probable non-vertebrate infecting” and excluded from further analysis. No vertebrate-like candidates were found in *Appendicularia* libraries and only three candidates were detected among the *Thaliacea* libraries (**Fig. 1d**).

Because each viral taxon with known vertebrate hosts also includes families of viruses that infect non-vertebrates (i.e., invertebrates and organisms outside of the animal kingdom), we performed preliminary phylogenetic analyses to identify and remove vertebrate-like candidates that fell within known invertebrate-infecting virus clades. Virus candidates belonging to the orders *Amarillovirales*, *Birnaviridae*, *Bunyavirales*, *Hepelivirales*, *Nodamuvirales*, *Picornavirales*, and *Reovirales* shared closer phylogenetic relationships to non-vertebrate-associated viruses in their respective taxa (**Figs. S1-9**). Notably, two divergent *Amarillovirales* NS5 sequences were positioned close to Chowder bay tunicate associated flavi-like virus^22^ near the tamanaviruses (**Fig. S1b**). Two tunicate-associated birna-like viruses formed a clade with Chicken proventricular necrosis virus and a bat faecal birnavirus (UVF62127.1) with low support when aligned with MAFFT (**Fig. S2a**), but because this relationship was not apparent when a different alignment method (MUSCLE^23^) was used (**Fig. S2b**), we were unable to draw a confident conclusion on the phylogenetic relationships among these viruses. Similarly, two viruses found in Stolidobranchia libraries (*Polycarpa*, accession: SRR18265385; *Botrylloides*, primary sample) fell at the base of a fish astrovirus clade (**Fig. S9a**). However, this relationship was not recapitulated when sequences were aligned with MUSCLE^23^. Instead, these viruses fell at the base of a clade comprising bat-associated astroviruses and Duck hepatitis virus 3 (albeit with low support; sh-aLRT/UFboot = 50/59) (**Fig. S9b**). In the case of the *Nidovirales*, tunicate-associated nidoviruses were not closely related to the vertebrate-infecting members of this order and exhibited unique features that we discuss below.

As a consequence of these analyses, we removed viruses belonging to the *Amarillovirales*, *Birnaviridae*, *Bunyavirales*, *Hepelivirales*, *Nodamuvirales*, *Picornavirales*, and *Reovirales* from our vertebrate-like candidate data set. Virus candidates belonging to the *Articulavirales, Jingchuvirales, Martellivirales, Mononegavirales,* and *Nidovirales* were retained for subsequent analysis.

### Evolutionary history of the influenza virus lineage may have predated the Vertebrata

We detected articulavirus-like segments in eight *Ascidiacea* libraries (**Fig. 2a**). Three libraries (accessions SRR7503161, SRR7503162, SRR7503170) were from the same BioProject (PRJNA480375) and their corresponding segments were identical. These and the library generated from *Clavelina lepadiformis* (accessions ERR11767553, ERR11767551) were derived from pharyngeal tissue. Only a small fragment (692nt) of a polymerase basic (PB) 1-like segment was found in a primary library that we collected at the first time point. Two fragments of a PB3-like segment were also identified in this library but could not be joined with PCR. We identified complete or near-complete additional segments in five of the eight libraries and fragments in two libraries (**Fig. 2a**). All segments were at low abundance in their respective libraries (<0.01% non-rRNA reads) (**Supp. Data 1**).

**Figure 2.**
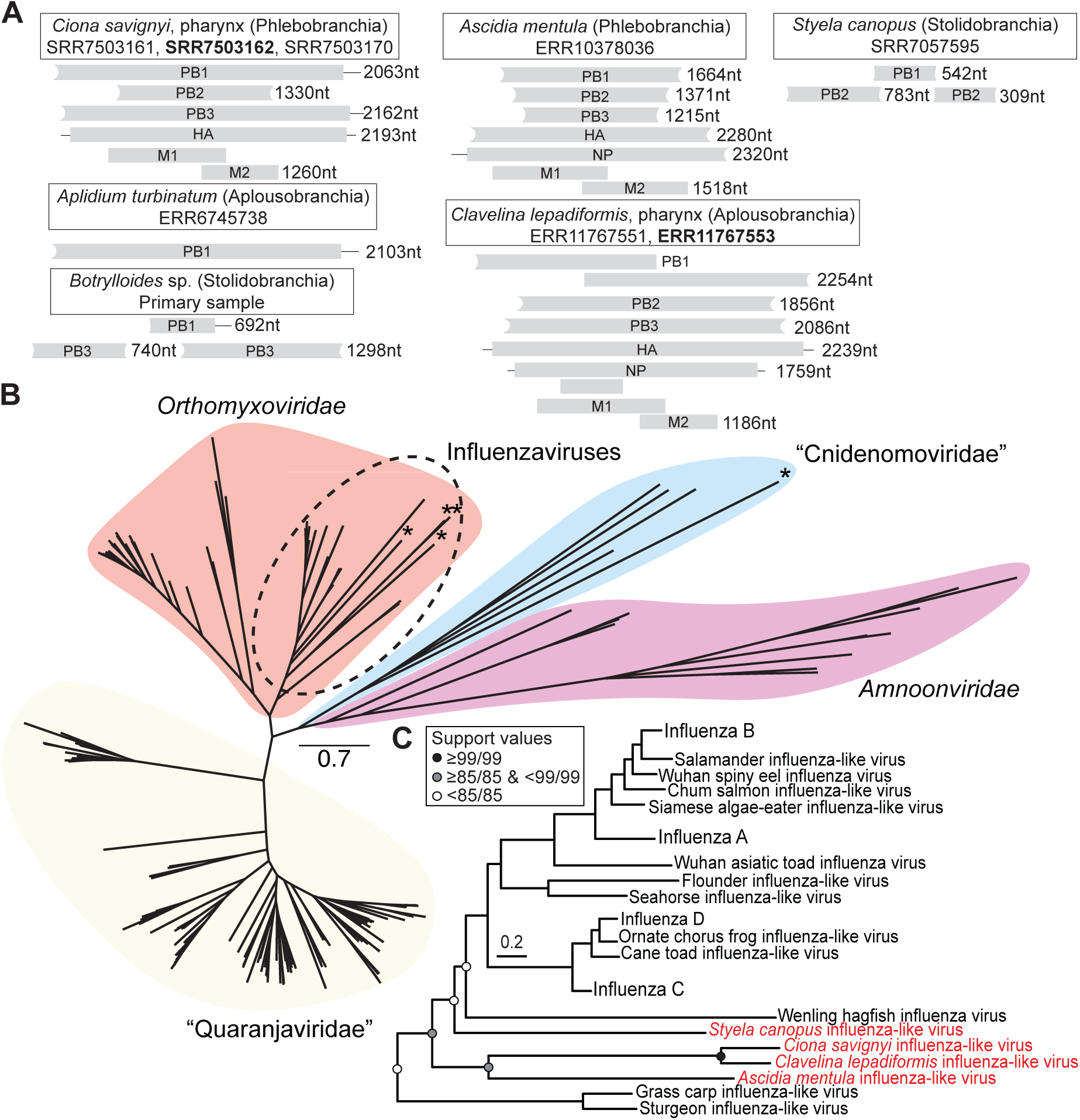
Influenza-like virus segments in seven ascidian metatranscriptomes. (a) Schematic of articulavirus-like segments detected in SRA (n = 8) and primary (n = 1) metatranscriptome libraries. The lengths of each segment are denoted in nucleotides (nt). Curved edges indicate incomplete ORFs. Bold face indicates the representative library from which segments were visualised. (b) Unrooted phylogeny of the *Articulavirales* inferred using the PB1 gene. Viruses identified in this study are indicated with stars. (c) Extract of the influenza virus clade from the order-level phylogeny. Viruses identified in this study are denoted by red tip labels. Support values for select nodes are reported as sh-aLRT/UFboot. The branches of both phylogenies are scaled by the number of amino acid substitutions per site.

Notably, the PB1 segment in the *Clavelina* library encoded a complete PB1 protein with two overlapping ORFs that could make the complete PB1 through a programmed ribosomal frameshift (-1 PRS). This -1 PRF mechanism involves a “slippery” heptanucleotide sequence (X_XXY_YYZ, here A_AAA_AAA), followed by a downstream stem-loop RNA structure, stimulating ribosomal slippage. The placement of this frameshift (position 841) was not consistent with the position of the PB1-F2 ORF found in some influenza A PB1 segments^24^. The sequencing coverage for this contig was low (median fold = 6, BBMap, **Fig. S10**), but it was not indicative of an assembly error. We detected five additional segments in this library (PB2, PB3, NP, HA, M), which supported the conclusion that this virus was exogenous rather than an endogenous sequence.

Four of the articulavirus-like PB1 segments were influenza-like, falling basal to the hagfish-associated PB1-like sequence with long branch lengths (**Fig. 2b,c**). Three of these (*Ciona savignyi*, *Clavelina lepadiformis*, and *Ascidia mentula* influenza-like viruses) formed a clade (**Fig. 2b,c**). Although the relationship of these sequences to that associated with a hagfish (jawless vertebrate) and most other vertebrate-associated influenza viruses was consistent with virus-host co-divergence over timescales of hundreds of millions of years (**Fig. 2c**), their relationship to each other was not. In particular, *Ciona* and *Ascidia* are both members of the Phlebobranchia, while *Clavelina*, the sister group to the *Ciona*-associated sequence, belongs to the *Aplousobranchia*. Additionally, two fish-associated sequences (Grass carp influenza-like virus and Sturgeon influenza-like virus) fell at the base of the clade, albeit with low support (sh-aLRT/UFboot = 49.9/44), suggestive of possible ancient host jumps. This placement was consistent in the other two polymerase segments (PB2 and PB3, **Fig. S11**), but not in the structural proteins (HA and NP, **Fig. S12,13**) that may be too divergent to compare.

The other segments from the tunicate libraries similarly shared a close relationship with the canonical influenza viruses (i.e., Influenza A, B, C, and D) (**Fig. S11-13**). In all cases, the PB2 and PB3 segments shared a closer relationship with influenza C and D viruses rather than falling as distinct lineages (**Fig. S11**). The placement of the NP and HA segments was uncertain and changed depending on the alignment method used (MAFFT: C/D-like, MUSCLE: A/B-like) (**Fig. S12-13**). The PB1 segment from the *Aplidium* library (ERR6745738), which was the only segment we could identify in this library, was not influenza-like and formed a sister group with the divergent “Cnidenomoviridae”^21^ (**Fig. 2a,b**).

Given that influenza-like viruses have not been previously documented in invertebrates and ascidians are filter-feeders that consume particulate organic matter, an alternative hypothesis was that these viruses were derived from the diet of the host animal. However, this was not supported through an analysis of the library composition. In three cases, other chordate reads comprised a small percentage of each library (6% ERR1037086, 1% SRR7503170, 0.7% Primary Sample #17), and no other chordate reads were detected in the remaining four libraries (**Fig. S14**). The consistently close phylogenetic relationship of the tunicate-associated segments further suggested that tunicates were the *bona fide* host because the libraries were derived from organisms collected across multiple continents (e.g., ERR10378036, Europe; SRR7503170, Asia; Primary sample #17, Oceania). We therefore conclude that these viruses likely represent the first observed instances of invertebrate-associated influenza-like viruses, and, as a consequence, that the influenza virus lineage evolved prior to the Vertebrata.

### Evidence for aquatic, invertebrate origins of multiple vertebrate disease-causing viral lineages

Given the discovery of tunicate-associated influenza-like viruses we asked whether other disease-causing viral lineages might have originated early in chordate evolution. We identified one such example in the novirhabdoviruses, negative-sense RNA viruses from the order *Mononegavirales* that can cause overt disease in finfish^25^. Specifically, five putative viral contigs in our data set formed a “stepwise” branching pattern at the base of the canonical *Novirhabdovirus* clade (**Fig. 3a**). The family-level tree topology recapitulated the position of the novirhabdoviruses as the only members of the *Gammarhabdovirinae*, which form a sister group to the other lineages within the *Rhabodiviridae*^26^ (**Fig. 3b**). Two novirhabdo-like viruses were associated with Stolidobranchia (*Botryllus schlosseri*, accession SRR10352255; *Botrylloides leachii*, primary sample). Three identical and highly divergent Phlebobranchia-associated viruses from the same BioProject (PRJNA248170, *Ciona intestinalis*) fell between them (**Fig. 3b**). The *Ciona* libraries were derived from nervous tissue in animals undergoing regeneration (**Fig. S15**). Two novirhabdo-like contigs encoded seemingly complete genomes (13306nt) with six ORFs, while the third contig was partial (8764nt) (**Fig. 3b**).

**Figure 3.**
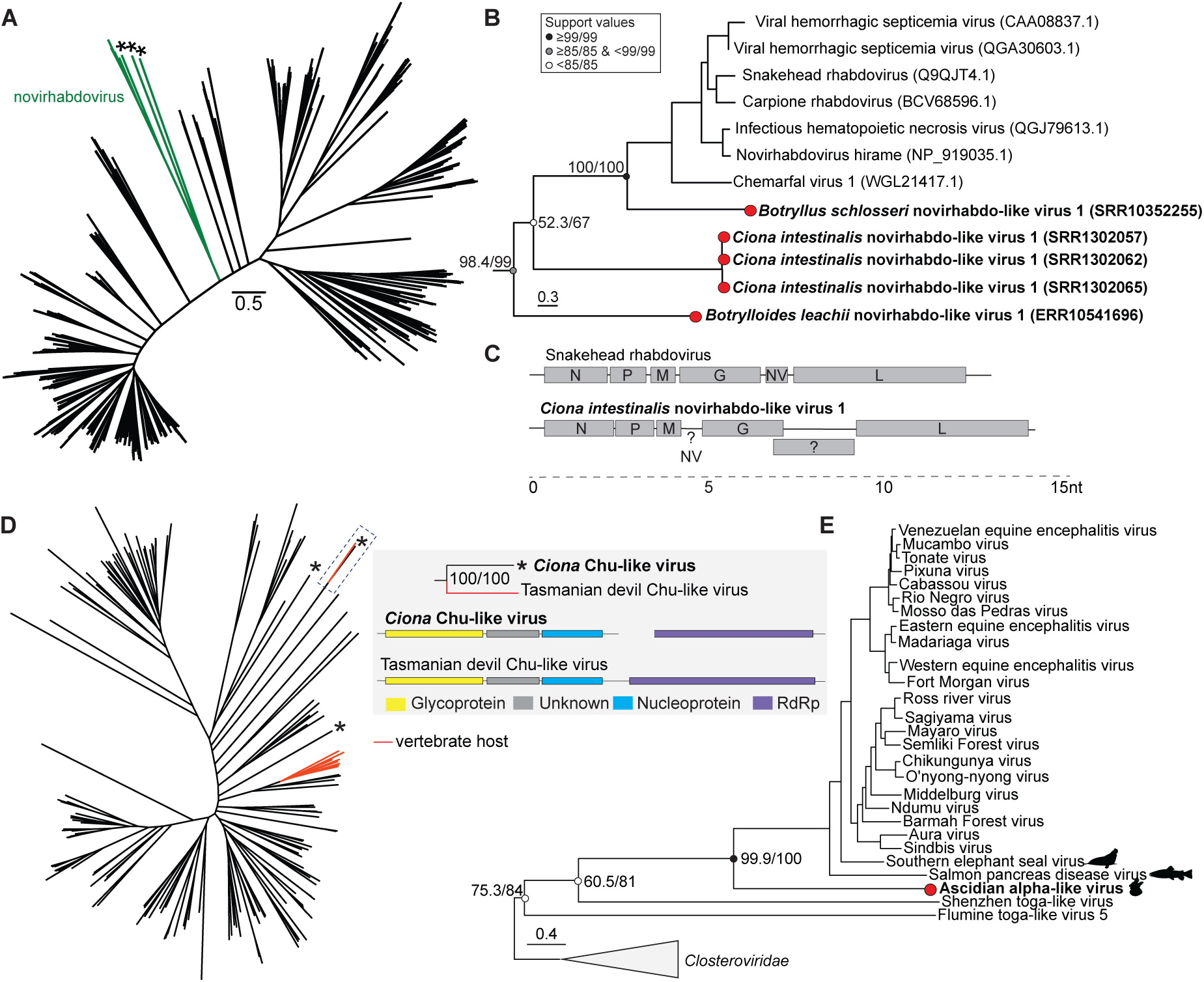
Evolutionary origin of the novirhabdoviruses, vertebrate-infecting *Jingchuvirales*, and the *Togaviridae* may have predated Vertebrata. For all phylogenies, branches are scaled according to the number of amino acid substitutions per site. (a) Phylogenetic diversity of the *Rhabdoviridae* with the novirhabdoviruses indicated by green branches. Stars denote viruses identified in this study. (b) The *Novirhabdovirus* clade from (a). Support values are shown as sh-aLRT/UFboot at select nodes. Red tip circles denote viruses identified in this study. (c) Genomic organisation of a canonical novirhabdovirus (Snakehead rhabdovirus) and a novirhabdo-like virus identified in this study (*Ciona intestinalis* novirhabdo-like virus 1). Genomes are scaled by nucleotides (nt). (d) Phylogenetic diversity of the order *Jingchuvirales* RdRp-encoding ORFs. Viruses that are associated with vertebrate hosts are indicated by red branches. Stars denote viruses identified in this study. Inset: Genomic organisation of *Ciona* Chu-like virus identified in this study and Tasmanian devil Chu-like virus. (e) Rooted phylogeny of the *Togaviridae* (order *Martellivirales*). Support values are shown as sh-aLRT/UFboot at select nodes. The red circle indicates the placement of the ascidian-associated alpha-like virus identified in this study. The icons represent the putative host of select viruses (source: phylopic.org).

The genomic architecture of the seemingly complete novirhabdo-like virus differed from that of canonical novirhabdoviruses. Specifically, the *Ciona*-associated virus (*Ciona intestinalis* novirhabdo-like virus 1) was ∼200nt longer than other viruses in this genus, and the ORF encoding the glycoprotein appeared to be split with a frameshift (**Fig. 3c**). In addition, we were unable to identify the non-virion (NV) protein unique to the novirhabdoviruses using sequence-based similarity methods. It may be that the tunicate-associated NV is too divergent in sequence, and we hypothesised that it could be encoded between ORFs 3 and 4 (**Fig. 3c**), although it is also plausible that these viruses do not encode a NV protein at all. We therefore concluded that these ascidian-associated viruses were novirhabdo-like and represented divergent, invertebrate members of the genus.

In a similar fashion, we detected two fragments of a chu-like virus closely related to Tasmanian devil chu-like virus (TDCV)^27^ in a library generated from *Ciona intestinalis* larvae undergoing tail regression (accession SRR18365523, BioProject PRJNA725676). We assumed that the ascidian was the host because no other plausible host was present in the library, which comprised >99% Phlebobranchia contigs. One fragment encoded the RNA-dependent RNA polymerase (RdRp) and was present at low abundance (0.003% of non-rRNA reads). The 5’ terminal of the ORF was incomplete, precluding us from determining whether this virus is segmented. Nevertheless, the RdRp sequence formed a sister clade with that of the TDCV (sh-aLRT/UFboot = 100/100), and this clade did not fall within any known *Jingchuvirales* lineages (**Fig. 3d, Fig. S16**) as previously reported^28^. The other fragment encoded three ORFs (a metapneumo-like glycoprotein, a protein of unknown function, and a bornavirus-like nucleoprotein), which reflected the genome organisation of TDCV (**Fig. 3d, *inset***) and was more abundant than the RdRp-encoding fragment (0.13% non-rRNA reads). The nucleoprotein and the glycoprotein sequences similarly formed clades with their TDCV counterparts (**Fig. S17**). Taken together, these evolutionary relationships, the shared genome organisation, and the phylogenetic distance from other vertebrate-infecting viruses in this order, suggested that there are multiple unrelated clades of chordate-infecting viruses in the *Jingchuvirales*.

Lastly, we identified a fragment (3481nt) of an alpha-like virus in a library generated from a hemolymph sample of an *Ascidia sydneiensis samea* (Phlebobranchia) at low abundance (0.002% of non-rRNA reads). Although the ORFs encoding non-structural and structural proteins were both partial, they exhibited well-defined alphavirus characteristics. The structural ORF encoded an alphavirus capsid protein (HHPred probability = 97.16%, e-value = 0.00027). The RdRp encoded by the non-structural ORF fell at the base of the canonical alphavirus clade with robust support (sh-aLRT/UFboot = 99.9/100) (**Fig. 3e, Fig. S18**). Of note, its relationship basal to the two known aquatic alphaviruses (southern elephant seal virus and salmon pancreas disease virus, **Fig. 3e, *icons***) was consistent with virus-host co-divergence and an aquatic origin of the *Togaviridae*^29^.

### Recombinant history of ascidian-associated nidovirus glycoproteins

Thus far, we have examined the evolutionary history of RNA viral taxa using the RdRp. However, other viral proteins, such as those that facilitate virus entry into the host cell, can exhibit markedly different histories^30^. Recombination involving glycoproteins has already been reported in tunicate-associated nidoviruses. Specifically, two of these viruses belong to the *Medioniviridae* (positive-sense RNA viruses; order *Nidovirales*) and encode Class II fusion proteins with apparent similarity to togavirus E1 glycoproteins (*Botrylloides leachii* nidovirus, MK956105 and Tognidovirus botryllus22085, BK066634^19^) – hence the name “Tognidoviruses”. This contrasts with the glycoproteins encoded by other chordate- and invertebrate-infecting families in the order: the *Coronaviridae*^31^, *Tobaniviridae*^32^, and *Mesoniviridae*^33^ that utilise Class I fusion proteins, and the *Arteriviridae*, whose fusion protein has yet to be identified^34,35^. Thus, recombination among diverse virus groups may have played a central role in shaping the evolution of the *Nidovirales* within chordates.

To investigate recombination in more detail, we first identified six additional tunicate-associated nido-like viruses. Two of these – an apparently complete 28kb putative viral contig in a *Botrylloides* ascidian and a partial RdRp fragment in a *Didemnum* ascidian – shared sequence similarity with members of the *Medioniviridae* (**Fig. 4a**). The remaining four were highly divergent and present in 53 libraries (51 *Phlebobranchia*, 2 *Aplousobranchia*) (**Supp. Data 2**). We recovered only fragments of two of the four viruses, but the other two appeared to be complete and were ∼33kb and ∼40kb in length. They did not encode a frameshift in their putative ORF1ab and expressed either one or two ORFs following the putative glycoprotein (**Fig. 4a**). We assigned names to these viruses according to their putative hosts (**Fig. 4a**) and will use them herein.

**Figure 4.**
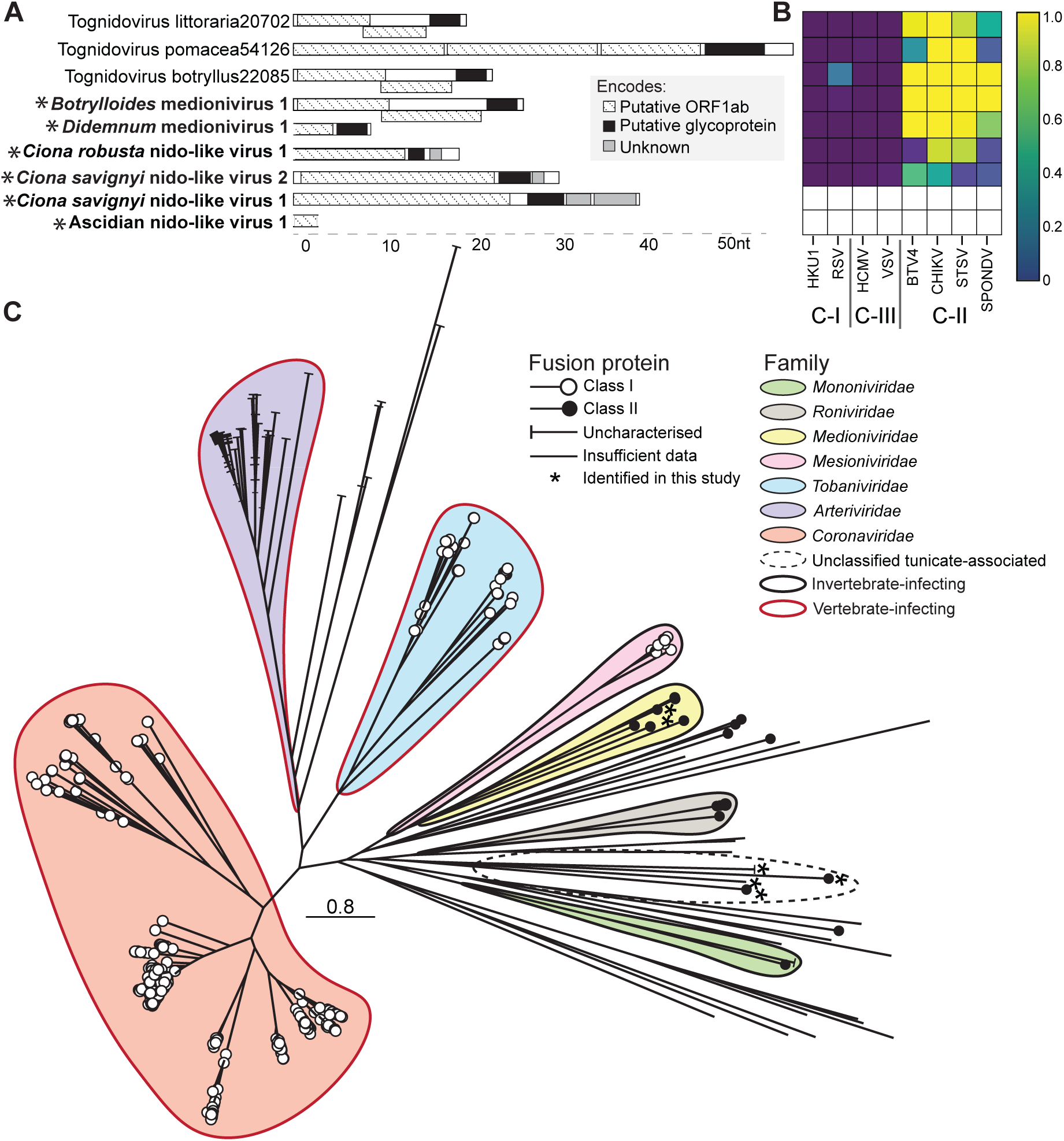
Utilisation of Class I and Class II fusion proteins across the *Nidovirales*. (a) Genomic architecture of tunicate-associated nido-like viruses and previously identified “Tognidoviruses”. Asterisks indicate viruses identified in this study. Genomes are scaled by genome size (nt). (b) Protein structure predictions for sequential overlapping sequence blocks, spanning entire viral polyproteins, were compared to class-I (C-I), III (C-III) and II (C-II) fusion protein reference structures using Foldseek^37^. Heat map displays highest detected Foldseek probabilities for each nido- and nido-like virus against each fusion protein reference. White squares are shown where no probability value was returned by Foldseek. (c) Unrooted phylogeny of the *Nidovirales* RdRp. Branches are scaled by the number of amino acid substitutions per site.

We next used protein structural prediction to identify and characterise the glycoproteins of these viruses and of members of nidovirus families for which the class of fusion protein was not known. Class II fusion proteins were a defining characteristic of the *Medioniviridae*, including the members we identified in this study (“*Botrylloides* medionivirus 1”, “*Didemnum* medionivirus 1”, **Fig 4b**). Of note, they shared nearly equal structural homology (defined by FoldSeek probabilities) with the glycoproteins of flaviviruses (Spondweni virus and Bole Tick virus 4), togaviruses (chikungunya virus), and bunyaviruses (Severe Fever with Thrombocytopenia Syndrome Virus), suggesting that their glycoproteins share a deep ancestry with all Class II fusion proteins and are not necessarily “toga-like” (**Fig 4b**). The glycoproteins of the non-medioniviruses identified here (“*Ciona robusta* nido-like virus 1”, “*Ciona savignyi* nido-like virus 1 & 2”) were more difficult to classify, with sequence-based analyses returning ambiguous results (i.e., low probabilities and high e-values, **Table 1**). However, by structure-based analysis, *Ciona robusta* nido-like virus 1 and *Ciona savignyi* nido-like virus 2 glycoproteins were inferred to be Class II-like with medium to high probability (0.912 and 0.692, respectively, **Fig 4b**). The protein structure of the *Ciona savignyi* nido-like virus 1 putative glycoprotein could not be predicted confidently, suggesting it is highly divergent.

**Table 1.** HHPred results for glycoproteins encoded by divergent ascidian-associated nidoviruses.

| Virus name | HHPred reference name | E-value | Probability (%) |
| --- | --- | --- | --- |
| *Ciona savignyi* nido-like virus 1 | Envelope gB; Varicella Zoster Virus | 4.2 | 67.5 |
| *Ciona savignyi* nido-like virus 2 | Glycoprotein, zika virus | 5.1 | 84.85 |
| *Ciona robusta* nido-like virus 1 | Protocadherin-15 | 3.3 | 81.37 |

The glycoproteins of previously identified nidoviruses were also predominantly Class II-like. According to sequence-based analyses, the *Roniviridae* and *Euroniviridae* utilise fusion proteins with similarity to both alpha- and bunyavirus glycoproteins (**Table S1**). Unlike other members of the *Nidovirales*, the *Mononiviridae* encode a single polyprotein, and the location of the glycoprotein therein was previously unknown. We therefore searched for glycoprotein-like features in the Flatworm mononi-like virus polyprotein using protein structure prediction and found a fold (amino acid residues 9650-9800) that was consistent with the C-terminal portion of a Class II fusion protein (comprising Domain-III and the transmembrane region; **Fig. S19**). Similarly, divergent nidoviruses previously identified using artificial intelligence-based analyses^36^ encode Class II-like fusion proteins, albeit with varying degrees of likelihood (**Table S2, Fig. S20**).

Given the frequent occurrence of Class II fusion proteins among nidovirus families, we mapped fusion protein class onto the *Nidovirales* RdRp phylogeny to assess the frequency of inter-virus recombination. Notably, fusion protein types fell on multiple branches on the RdRp phylogenies, indicative of recombinant evolutionary histories. The two tunicate medioniviruses identified here fell within the diversity of the Class II-encoding *Medioniviridae,* which forms a sister group with the Class I-encoding *Mesioniviridae* (**Fig. 4c**). The remaining four newly identified tunicate nido-like viruses did not fall within a known family, and the long branches characterising the latter combined with their range of genome size (between ∼33kb and 40kb) suggest that each may represent novel virus families (**Fig. 4c**). Although there was an apparent division of vertebrate- and invertebrate-infecting lineages, this pattern again did not correspond to glycoprotein class. Thus, as with the flaviviruses^30^, glycoprotein composition in the *Nidovirales* is not congruent with the evolutionary history of the RdRp, such that recombination likely played a substantial role in shaping this order.

### Virus-host co-divergence and recombination in a divergent mononega-like virus

The examination of Class II fusion proteins in our tunicate virus data set revealed an additional example of inter-virus recombination of a glycoprotein in a highly divergent virus found in the *Styela rustica* hemolymph library (accession SRR16495746). The organisation of this putative virus resembled that of members of the *Rhabdoviridae*, a family of negative-sense RNA viruses (order *Mononegavirales*), encoding six complete ORFs, including a nucleoprotein, a glycoprotein, and an RdRp that utilises the GDD amino acid motif in the palm C catalytic triad (**Fig. 5a**). The first ORF was mononega-like, sharing similarity with *Borna-* and *Paramyxoviridae* nucleoproteins (**Table 3**). Although the length and position of ORFs 2 and 3 may correspond to rhabdo-like phosphoprotein (P) and matrix protein (M), respectively, they shared neither sequence nor structural homology with known proteins (**Fig. 5a**). We were also unable to determine the possible function of the sixth ORF (**Supp. Data 3**). The RdRp was highly divergent, sharing minimal sequence similarity (28.57%) with members of this order (**Table 4**). Phylogenetic analysis of the *Negarnaviricota* placed this virus sequence at the base of the vertebrate-infecting *Paramyxoviridae* and *Pneumoviridae*, suggesting that the ancestors of these families evolved among invertebrate, chordate hosts (**Fig. 5b**). We named this putative virus *Styela rustica* patchyvirus 1.

**Figure 5.**
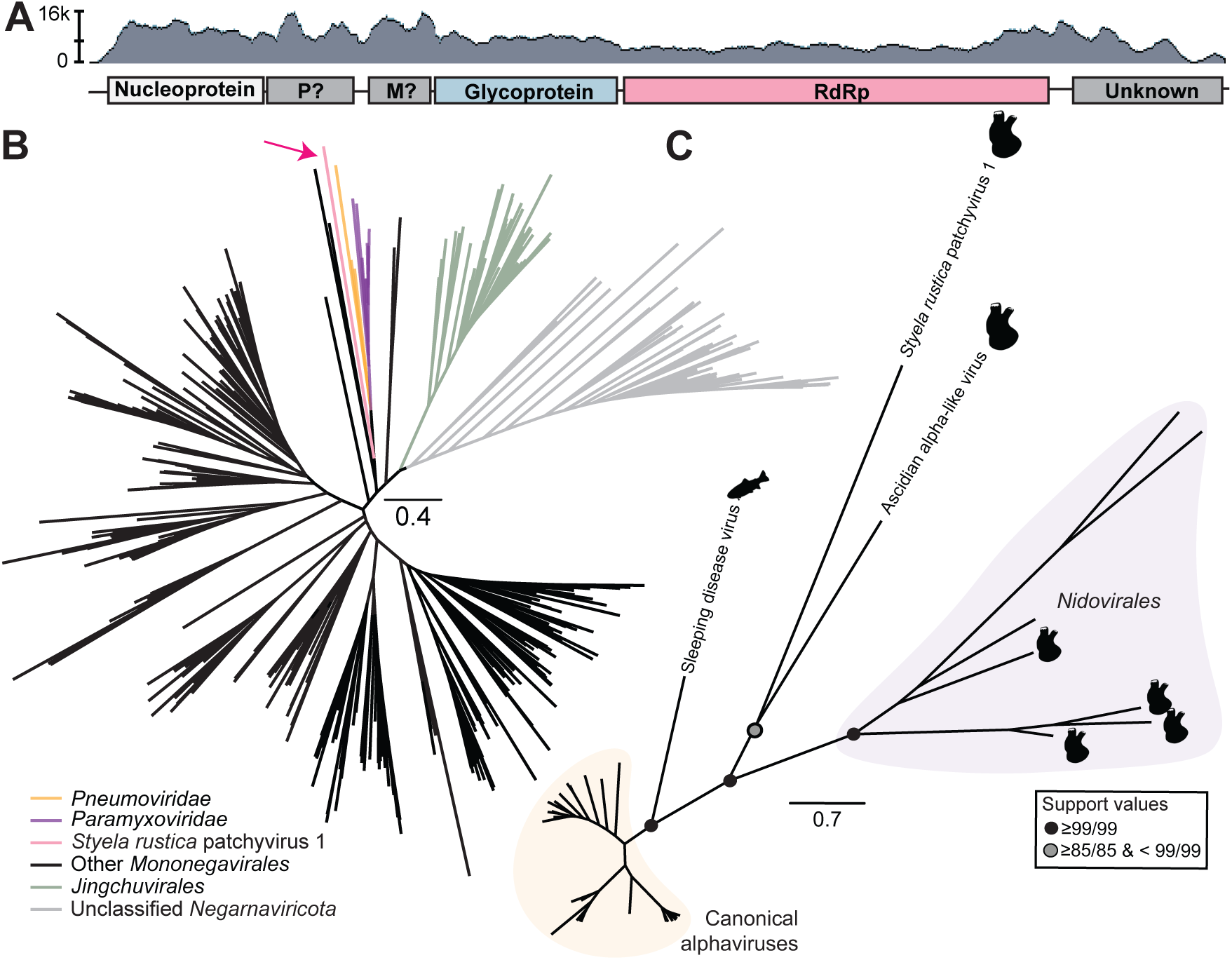
Recombination of Class II glycoproteins to a tunicate-associated mononega-like virus. (a) Schematic of divergent mononega-like viral genome sequence with sequencing coverage. Question marks indicate ORFs that shared no detectable sequence or structural similarity with known proteins but whose positions may be indicative of their function according to the genome organisation of the *Rhabdoviridae*. Abbreviations: P (phosphoprotein), M (matrix protein). (b) Unrooted maximum likelihood phylogeny of the *Negarnaviricota*. Branches are scaled by amino acid substitutions per site. The arrow indicates the placement of the *Styela rustica* patchyvirus 1 identified in this study. Branches are scaled by amino acid substitutions per site. (c) Unrooted maximum likelihood phylogeny of alphavirus E1 glycoproteins and class II glycoproteins encoded by tunicate-associated RNA viruses. Branches are scaled by amino acid substitutions per site, and support values are shown at select nodes as sh-aLRT/UFboot. Animal icons indicate the known or associated host of each putative virus (source: phylopic.org).

**Table 3.** HHPred results for the putative nucleoprotein encoded by Styela rustica patchyvirus 1.

| HHPred reference name | E-value | Probability (%) |
| --- | --- | --- |
| Borna disease virus nucleoprotein | 6.3e-33 | 100 |
| Newcastle disease virus nucleoprotein | 2.7e-6 | 98.6 |
| Parainfluenza virus 5 nucleocapsid | 3.3e-6 | 98.58 |
| Nipah virus nucleoprotein | 4.4e-6 | 98.53 |

**Table 4.** BLASTx nr results for Styela rustica patchyvirus 1 ORFs.

| Reference name (accession) | Query cover (%) | E-value | Identity (%) |
| --- | --- | --- | --- |
| *RdRp (ORF4)* |  |  |  |
| Beihai rhabdo-like virus 5 (YP_009333422.1) | 20 | 4e-04 | 28.57 |
| Hubei rhabdo-like virus 5 (YP_010084250.1) | 26 | 0.001 | 22.26 |
| *Artemisia capillaris* nucleorhabdovirus 1 (YP_010805763.1) | 29 | 0.002 | 23.59 |
| Jungle carpet python virus (YP_009508484.1) | 20 | 0.002 | 26.92 |
| *Glycoprotein (ORF3)* |  |  |  |
| Western equine encephalitis virus (ABD57956.1) | 49 | 3e-08 | 25.93 |
| Highlands J virus (ANJ02316.1) | 49 | 2e-06 | 25.83 |
| Fort Morgan virus (YP_003324599.1) | 59 | 3e-06 | 23.15 |
| Eastern equine encephalitis virus (ADB08674.1) | 55 | 6e-06 | 25.86 |

In a similar manner to the *Rhabdoviridae*, but unlike the *Paramyxoviridae* and *Pneumoviridae*, the genomic architecture of *Styela rustica* patchyvirus 1 suggested that it encoded a single glycoprotein ORF (**Fig. 5a**). Strikingly, this glycoprotein was not rhabdo-like (Class I), but instead shared low (25.93%) but detectable sequence similarity to the E1 glycoprotein of alphaviruses, a group of positive-sense RNA viruses (*Togaviridae)* (**Table 4**). When the phylogenetic relationship of all so-called “toga-like” glycoproteins was inferred relative to the canonical alphavirus glycoproteins, this sequence formed a sister group to the glycoprotein of Ascidian alpha-like virus described above, albeit with long branches (**Fig. 5c**). Due to the divergent nature of these sequences, we could not conclude that *Styela rustica* patchyvirus 1 acquired its Class II fusion protein from the *Togaviridae*; however, the shared sequence identity suggests that a recombination event between positive- and negative-sense RNA viruses may have occurred during the evolutionary history of this viral lineage. Thus, given its unique composition, *Styela rustica* patchyvirus 1 may be the first representative of a distinct family within the *Mononegavirales* that exemplifies a complex, recombinant evolutionary history.

## DISCUSSION

Our analysis of tunicate metatranscriptomes sheds new light on the evolutionary history of animal RNA viruses. Specifically, we present evidence that five vertebrate-infecting viral lineages may have evolved prior to the origin of the Vertebrata. The phylogenetic placement of the RdRp of some tunicate-associated influenza-, novirhabdo-, alpha-, chu-, and mononega-like viruses was consistent with a process of virus-host co-divergence that commenced in basal chordates. Perhaps most notable among these were the influenza viruses, which we previously hypothesised first emerged in early vertebrates due to basal fish-associated lineages^21^. In contrast, other tunicate-associated viruses belonging to nine viral lineages (the *Amarillovirales*, *Birnaviridae*, *Bunyavirales*, *Hepelivirales*, *Nidovirales*, *Nodamuvirales*, *Picornavirales*, *Reovirales*, and *Stellavirales*) were invertebrate-like or fell with non-animal infecting viruses. Whether this pattern reflects the evolutionary histories of those orders or is the result of insufficient sampling requires further investigation.

In addition to enriching our understanding of the origins of some disease-causing viral lineages, vertebrate-like tunicate-associated viruses exhibited unique genetic features. *Ciona* Chu-like virus and its closest relative Tasmanian devil Chu-like virus encode an ORF of unknown function between their chu-like glyco- and nucleoproteins. Similarly, *Ciona intestinalis* novirhabdo-like virus 1 may not encode the auxiliary non-virion (NV) protein. It could be that this protein, which has been shown to interfere with the host immune response^38^, was an innovation that evolved when the novirhabdoviruses emerged in vertebrates or that the tunicate-associated homolog is too divergent to detect with sequence similarity-based methods. This protein or its derivative could be encoded by the second glycoprotein-like domain immediately preceding the RdRp, congruent with the canonical novirhabdovirus organisation. Characterising these tunicate virus domains in cell culture would directly address the question of how the evolution of the vertebrate immune system may have influenced that of RNA viruses.

More striking was the example of *Styela rustica* patchyvirus 1, which comprises a negative-sense mononega-like nucleoprotein and RdRp, coupled with a Class II fusion protein that shares detectable sequence similarity to the E1 glycoprotein of a positive-sense alphavirus. The placement of the RdRp at the base of the *Paramyxoviridae* and *Pneumoviridae* is indicative of a viral lineage that extends back to the basal chordates. Both of those vertebrate-infecting families encode Class I fusion proteins, which suggests that the switch from Class II to Class I formed part of the transition from invertebrate chordate to vertebrate hosts. A similar pattern of major changes of glycoprotein composition concordant with the use of vertebrate hosts has been observed in members of the *Flaviviridae*^30^. However, the retention of sequence similarity to alphavirus E1 in the patchyvirus is puzzling, particularly because this degree of similarity was not retained by the putative glycoprotein encoded by Ascidian alpha-like virus. It is also intriguing because it suggests a recombination event between positive- and negative-sense viruses that may have occurred more recently than the transition to vertebrate hosts. Calibrating the timing and direction of this event will require additional sampling of the diversity of the patchyvirus lineage.

Despite these uncertainties, incongruence between the glycoprotein and RdRp phylogenies can be observed multiple times in tunicate-associated viruses, and we found evidence that this pattern is not specific to tunicate hosts. Tunicate-associated nidoviruses utilise an array of Class II-like fusion proteins that differ markedly from both each other and those used by other families in the order. Similarly, glycoproteins encoded by the non-chordate-infecting families the *Roniviridae, Euroniviridae,* and *Mononiviridae* share sequence and/or structural homology with Class II fusion proteins. The *Medioniviridae*, which comprise a combination of tunicate and non-chordate invertebrate hosts, also exhibit this feature. In contrast, Class I fusion proteins are only utilised by the *Coronaviridae*, *Tobaniviridae*, and *Mesioniviridae* within the *Nidovirales*, and the RdRps of these families are polyphyletic. Unlike in the *Flaviviridae*, fusion protein type does not map onto the host: not all vertebrate-infecting families utilise Class I (e.g., the *Arteriviridae*) and not all invertebrate-infecting families encode Class II (e.g., the *Mesioniviridae*). Thus, host type does not account for such flexibility, and additional sampling is required to elucidate the evolutionary drivers that underpinned these recombination events.

Genome size also appears to be fluid within the *Nidovirales*. The discovery of Nam Dinh virus (family *Mesioniviridae*) prompted the hypothesis that this family represented an “evolutionary intermediate” between the large nidoviruses (the *Coronaviridae* and *Roniviridae*) and the small nidoviruses (the *Arteriviridae*), bolstering the argument that the acquisition of a proof-reading mechanism enabled the size of the nidovirus genome to expand^33^. The nido-like viruses discovered in this study and previously^19,36^ refute such a history. In particular, *Ciona savignyi* nido-like virus 1 (∼40kb) and *Ciona savignyi* nido-like virus 2 (∼30kb) formed a clade outside of any known family in the order. Not only is this relationship indicative of unrealised viral diversity, but it suggests that the genome size of nidoviruses has fluctuated, growing both larger and becoming smaller, at multiple points throughout the evolution of this order. Whether this is due to the acquisition of proof-reading mechanisms, to patterns of horizontal gene transfer or gene loss, or to other key genetic features requires experimental validation. It also raises the question of whether other orders of RNA viruses are equally disposed to take up genetic material from cellular organisms in this way. A large flavi-like virus that we previously identified in a sea sponge^39^, notable for both its length and apparent acquisition of bacterial genes, remains the sole example outside of the *Nidovirales*. Further exploration of the marine invertebrate virosphere is likely to reveal additional examples.

This study has some limitations. A primary challenge of metatranscriptomic analysis in the absence of experimental validation is the inability to directly determine the host associations of the newly identified viruses. We were instead reliant on indirect lines of evidence such as the library composition and the phylogenetic relationship of divergent viruses to one another to conclude that some of the viruses described in this study were likely infecting ascidians. Our data set was also affected by sampling biases, specifically the overrepresentation of *Ciona spp.* in the SRA. We were therefore unable to fully explore the viromes of the *Thaliacea* and *Appendicularia*, and a concerted effort to sample these organisms from their natural environments is needed to rectify these gaps. Additionally, although we could conclude that some viral lineages likely emerged before the origin of the vertebrates, we were unable to determine the timing of these events as we did not observe clustering by host order. For example, *Ciona intestinalis* novirhabdo-like virus 1 (host: *Phlebobranchia*) fell between two *Stolidobranchia*-associated viruses. A similar absence of clustering was observed for the influenza-like viruses, and the placement of the sturgeon- and carp-associated clade at the base of this group added another layer of virus-host incongruence. A further complication is a lack of consensus on the phylogenetic relationships within the Tunicata and even the *Ascidiacea*^40,41^, such that whether or not tunicate-associated viruses adhere to virus-host co-divergence within this subphylum cannot be assessed currently.

Despite this, our findings address and raise intriguing questions about the mechanisms and antiquity of animal RNA virus evolution. Virus-host co-divergence has, in multiple instances, been punctuated by the acquisition of genetic material through recombination across virus taxa that may have had a marked effect on virus evolution (e.g., *Styela rustica* patchyvirus 1). As this was the first systematic analysis of tunicate-associated RNA viruses, more systematic sampling of the Tunicata may find that the origins of additional disease-causing vertebrate viruses predated the Vertebrata. In particular, mapping the acquisition and loss of genetic features to the transition from basal chordates to vertebrates in these lineages could reveal viral traits that are correlated with disease emergence in vertebrates. Not only are tunicates a rich source of RNA virus diversity, but their viruses stand to further reshape our understanding of animal RNA virus evolution.

## METHODS

### Sample collection

Primary samples were collected as previously described^39^ from Chowder Bay, Sydney Harbour, Australia on November 24, 2021 (n = 12) and October 20, 2022 (n = 18). To minimise the risk of cross-contamination, divers wore latex gloves and cleaned forceps with 96% ethanol between sampling events. Samples were stored in RNA/DNA-free cryogenic tubes and placed immediately in liquid Nitrogen before being transferred to a -80C freezer for long-term storage.

### Classification of primary samples

All ascidians were identified based on morphology. Further classification was performed in two ways. The composition of each library was first analysed using CCMetagen^42^, KMA^43^, and and the NCBI nt database (June 2019). In many cases, this proved inconclusive due to the complex nature of the libraries, which include the ascidian, its diet, and the invertebrate symbionts that can inhabit the organism (**Fig. S21**). Second, the rRNA in each library was assembled and screened against the NCBI nucleotide (nt) database using blast+ v.2.12.0^44^ (**Supp. Data**).

### RNA extractions and sequencing

To extract total RNA, samples were processed individually and homogenised by flash freezing with liquid nitrogen and grinding with a sterile mortar and pestle (one mortar/pestle pair per sample). Total RNA was extracted using the QIAGEN RNeasy Plus Mini kit, with the RNA from some libraries pooled for downstream processing. Samples from the first time point were sequenced individually, and samples from the second time point were combined into eleven pools. Library preparation was performed using the Ribo-Zero Plus library preparation kit, and all libraries were sequenced on the Illumina NovaSeq 6000 platform. One “blank” negative control that contained sterile water and reagent mix was included for each run.

### Library processing and assembly

Nextera adapters were removed from raw reads using cutadapt v1.8.3^45^ and the following parameters: minimum length = 25, quality-cutoff = 24, number of bases trimmed at the beginning and end of reads = 5. The quality of trimming was checked FastQC v0.11.8^46^. In cases where removal was incomplete, the contigs were manually trimmed and reassembled if needed. Reads mapping to rRNA were removed using sortMeRNA v4.3.3^47^. Contigs were assembled with MEGAHIT v1.2.9^48^. A total of 3,468 libraries were successfully assembled (**Supp. Data 4**).

### SRA screening

To capture all publicly available tunicate metatranscriptomic data, we queried the NCBI SRA database for all libraries sequenced from “Tunicata”, “Urochordata”, “Sea squirt”, “Ascidiacea” as of March 2024, at least 0.5Gb in size (n = 3937). Ten libraries were removed because they were in fact associated with mollusc metatranscriptomes (1 Katharina tunicata, 1 Crobula tunicata, 8 Clione limacina).

### Identifying putative viral contigs

To detect RNA virus-like sequences, we screened all assembled libraries against the protein version of the Reference Viral Database (RVDB, November 2023)^49,50^ and RdRp-Scan databases using DIAMOND BLASTx v2.0.9^51^. We specified an e-value of 1 x 10^-5^ and the parameter “–very-sensitive”. All contigs that returned hits to viruses were checked against the NCBI non-redundant (nr) protein database (as of September 2024) and contigs that shared detectable sequence similarity to known cellular genes were removed from the data set. Minimum length thresholds for contigs were assigned according to order (**Table S2**).

All contigs were checked for open reading frames (ORFs) using the standard genetic code (all virus contigs), invertebrate mitochondrial genetic code (*Cryppavirales*), or ciliate genetic code (Zhaovirus). Contigs that encoded truncated ORFs or did not encode ORFs were removed from the data set. The sequencing coverage of contigs with unique features such as frameshifts or instances of putative recombination was assessed using BBMap v37.98^52^. Nucleotide sequences were translated using the Expasy web translate tool (https://web.expasy.org/translate/), EMBOSS getorf (https://www.bioinformatics.nl/cgi-bin/emboss/getorf), or InterProScan^53^. Abundance was measured according to the expected count generated by RSEM v1.3.0^54^.

### PCR of ascidian-associated PB3 fragments

PCR was performed using the SuperScript IV One-Step RT-PCR kit following the manufacturer’s instructions. An annealing temperature of 55C was used. Custom primers were designed (**Table 5**).

**Table 5.**
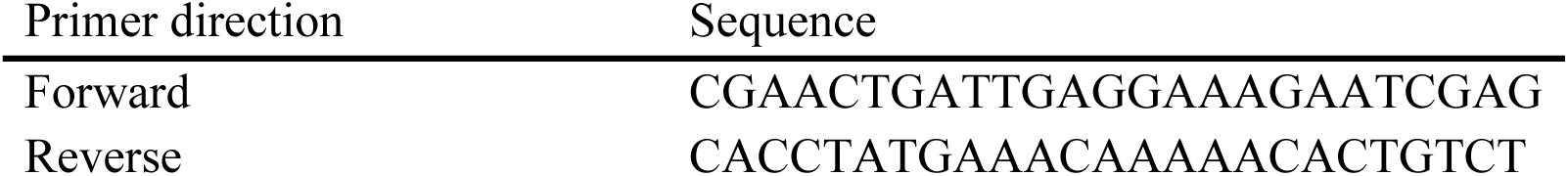
Primers used for RT-PCR.

### Genome annotation and protein structure prediction

We used multiple sequence-based methods to identify functional domains in the genomes of putative viral contigs: NCBI Conserved Domains^55^, InterProScan (CDD, NCBIfam, TMHMM)^53^, HHPred^56^, and Phyre2^57^.

We used protein structure prediction to search for putative glycoproteins as described previously^30,39^. Briefly, viral polyprotein sequences were split into sequential 300 residue blocks, overlapping by 100 residues. Structures were predicted using AlphaFold2-ColabFold and ESMFold^58–60^ and then compared to reference protein structures using Foldseek^37^. The following reference structures were used: Class I – HKU1 Spike (PDB: 5I08), RSV F (6APB). Class III: HCMV Gb (7KDP), VSV G (5I2S); Class II – CHIKV E1 (6NK7), SPONDV E (6ZQI), SFTSV Gc (8ILQ) and Bole Tick Virus 4 E glycoprotein identified in a previous study^30,61–67^. For each virus species, we identified structure blocks with the best Foldseek probabilities (highest values) and e-values (lowest values) against each of the reference structures.

### Phylogenetic analysis

#### Preliminary invertebrate/vertebrate analyses

We first curated reference data sets for RNA virus orders with known vertebrate and invertebrate hosts using the NCBI Virus resource. For the *Bunyavirales*, *Martellivirales*, and *Picornavirales*, we selected representative members of each family within these orders due to the high levels of sequence divergence. For the other viral orders, we downloaded all publicly available RdRp amino acid sequences and removed redundant sequences at 85% identity with CD-HIT v4.8.1^68^. The remaining sequences were aligned with MAFFT v7.490^69^, and sequences that did not align correctly were removed. Misaligned sequences were mostly misannotated genes (i.e., they were not RdRps). We aligned the *Bunyavirales*, *Martellivirales*, and *Nidovirales* using MAFFT and MUSCLE v5^23^. MUSCLE performed better for the *Bunyavirales* (i.e., it did not incorrectly pull fragmented sequences to the centre of the alignment), and hence this alignment was used for further analysis. All other phylogenetic trees were inferred from sequence alignments generated with MAFFT v7.490. We removed ambiguities at specified thresholds for the *Martellivirales* (70%) and *Nidovirales* (75%) and with the parameter gappyout in all other cases using trimAl v1.4.1^70^. Phylogenies were inferred using the maximum likelihood (ML) method in IQ-TREE v1.6.12^71^, with the ModelFinder limited to LG models with “-mset”. Node support was measured with 1000 ultrafast bootstraps (UFboot) and sh-aLRT.

#### Family and lineage-specific analyses

For the phylogenetic analysis of influenza virus segments (i.e., PB2, PB3, NP, HA), representative sequences from the canonical influenza viruses A-D were included, along with amphibian and fish-associated influenza-like viruses. Representative thogotoviruses and quaranjaviruses were included as outgroups. Sequences were aligned with both MAFFT v7.490 and MUSCLE v5. No ambiguities were removed, but the ends of the alignments were trimmed for PB2 and NP.

In the case of the *Togaviridae*, we downloaded all polymerase sequences from this family that were at least 400 amino acid residues in length (n = 1,041) from NCBI. Redundant sequences were removed at 90% identity with CD-HIT, leaving 45 unique sequences in our data set. We then added the first 100 BLASTx hits for Ascidian alpha-like virus and removed redundancies at 90% identity with CD-HIT. For the *Jingchuvirales* nucleoprotein and glycoprotein phylogenies, we downloaded all corresponding sequences available on NCBI and removed redundant sequences at 99% using CD-HIT. Sequences from each data set were aligned with MAFFT v7.490 and the edges of the alignments were manually trimmed. Ambiguities were removed with trimAl and the parameter gappyout, and ML phylogenies were then inferred with IQ-TREE v1.6.12 and the best-fitting model was chosen by ModelFinder. Similarly, we downloaded all alphavirus glycoproteins from NCBI (n = 2307) and removed redundancies at 85% with CD-HIT. The resulting phylogenetic trees were inferred as described for the *Jingchuvirales* glycoproteins. We compiled reference sequences from the *Negarnaviricota* from NCBI, the LucaProt database^36^, and studies identifying viruses in environmental samples^36,72–75^. Amino acid sequences were aligned with MAFFT v7.480 and ambiguities were removed with trimAl v1.2rev59 (parameters: -gt 0.8 -st 0.005 -cons 30). The phylogeny was inferred using the ML method available in PhyML v3.3.3^76^, and the model was limited to LG.

## RESOURCE AVAILABILITY

Nucleotide sequences, amino acid sequences, multiple sequence alignments, and tree files are publicly available at https://github.com/mary-petrone/tunicate_viruses_2024.

## ACKNOWLEDGMENTS

This work was funded by a National Health & Medical Research Council (NHMRC) Investigator grant (GNT2017197) to E.C.H., an Australian Research Council (ARC) Discovery Project grant (DP240101313) to E.C.H. and M.E.P., and by AIR@InnoHK administered by the Innovation and Technology Commission, Hong Kong Special Administrative Region, China to E.C.H. We thank David Grunde-McLaughlin for suggesting the name “*Styela rustica* patchyvirus” and Chris Cooney and Alex McGrath for helping with sample collection.

## AUTHOR CONTRIBUTIONS

M.E.P., E.M.M., and E.C.H. conceived of the study. E.M.M. and O.T. collected the primary samples. M.E.P., J.G., R.H.P., K.V.B., J.C.O.M., S.M., M.S., and O.T. performed the analyses.

M.E.P. and E.C.H. drafted the manuscript. All authors reviewed and edited the manuscript.

