## Supplemental Figures S1-S21 for "Tunicate metatranscriptomes reveal ancient virus-host co-divergence and inter-order recombination in the evolutionary history of disease-causing viruses"

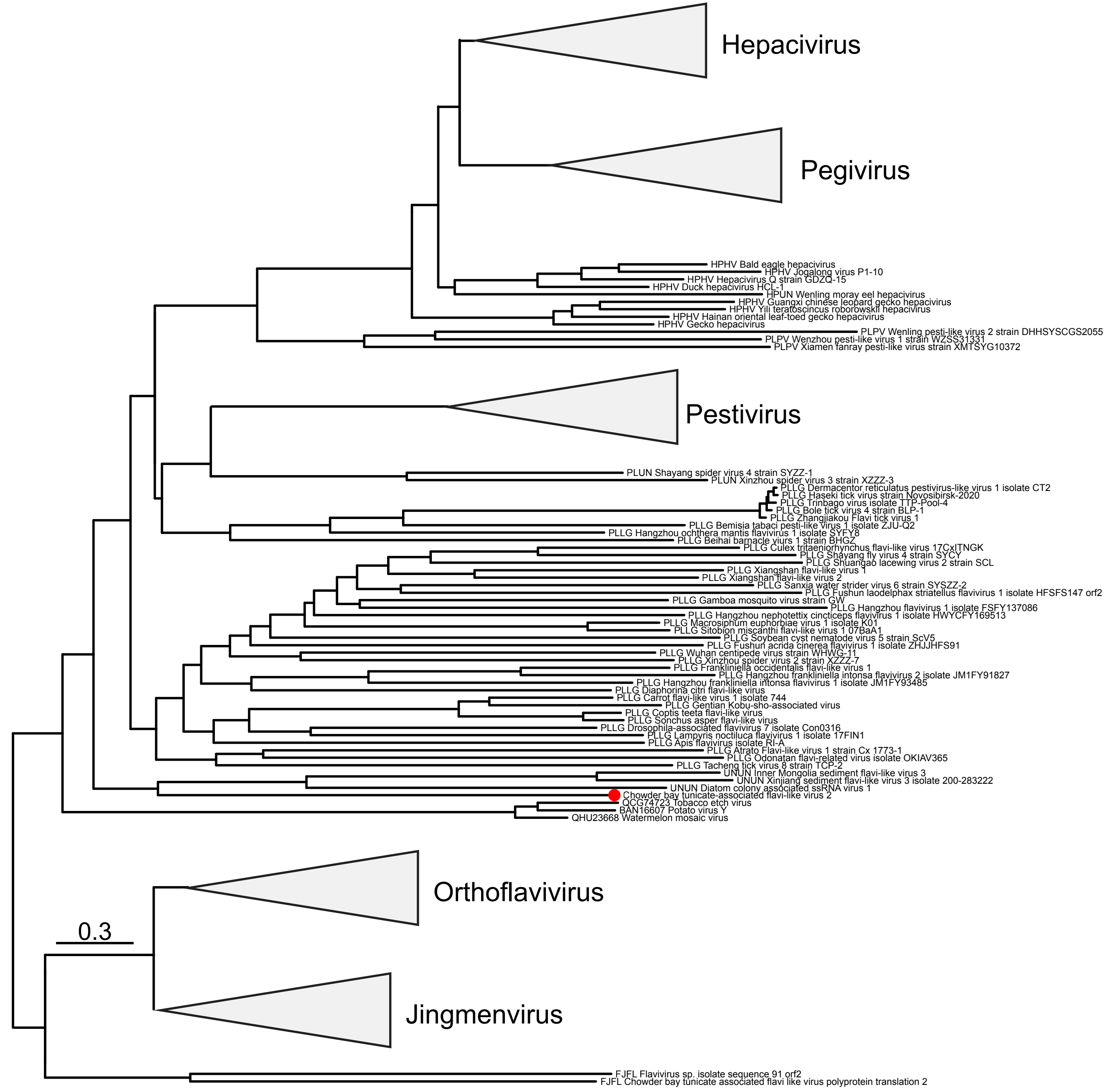


**Figure S1 Midpoint rooted phylogeny of the *Amarillovirales* NS3 gene.** Sequences in the tree were aligned with MAFFT v7.490, and the tree was inferred using IQ-TREE v1.6.12 with Model Finder limited to LG. Branches are scaled by amino acid substitutions. The red dot indicates the placement of the novel tunicate-associated virus fragmented identified in this study.

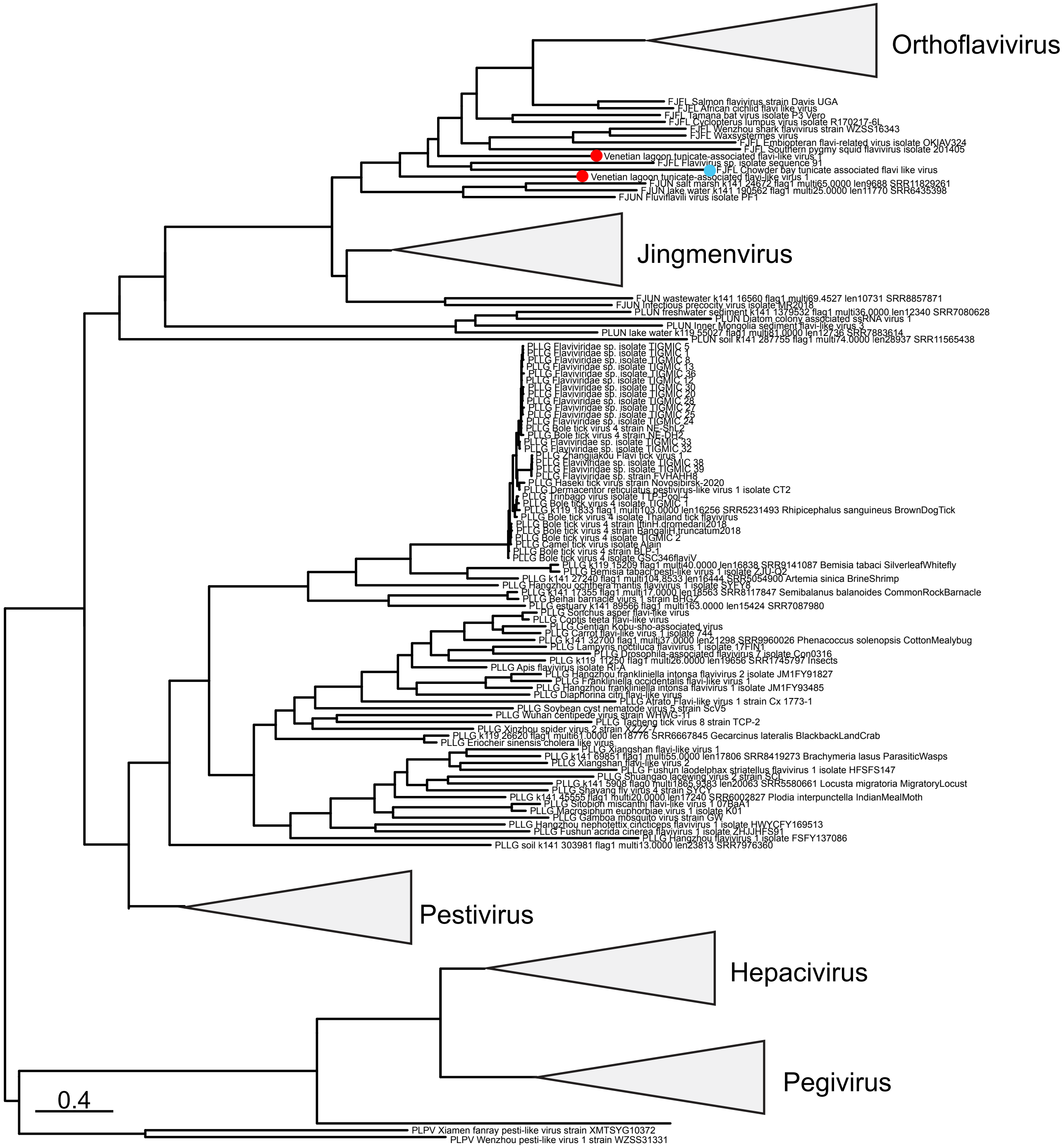


**Figure S2 Midpoint rooted phylogeny of the *Amarillovirales* NS5 gene.** Sequences in the tree were aligned with MAFFT v.7.490, and the tree was inferred using IQ-TREE v.1.6.12 with Model Finder limited to LG. Branches are scaled by amino acid substitutions. The red dots indicate the placement of the novel tunicate-associated virus fragmented identified in this study. The blue dot indicates the placement of Chowder bay tunicate-associated flavi-like virus.

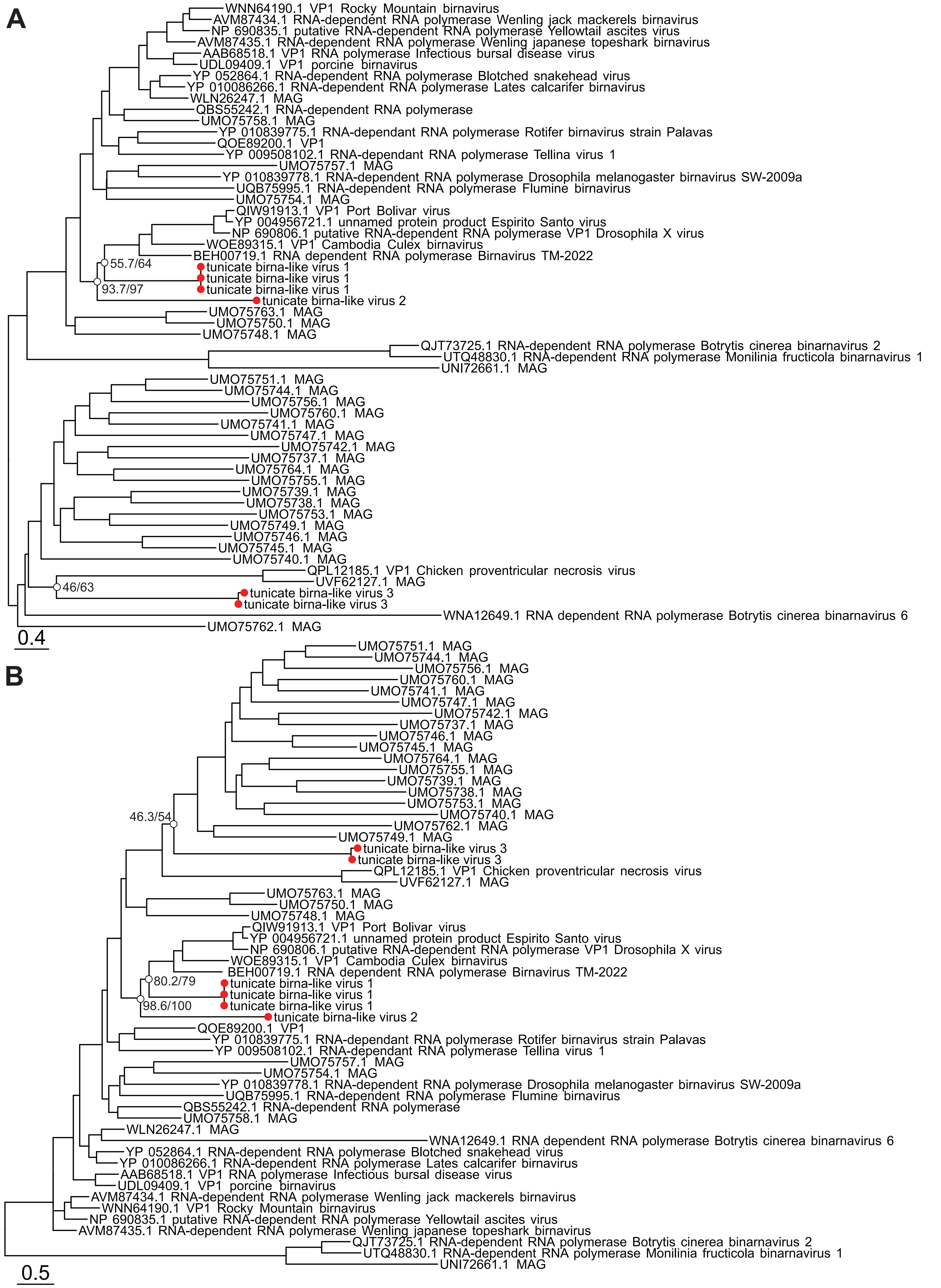


**Figure S3 Midpoint rooted phylogenies of the *Birnaviridae*.** (a) Sequences aligned with MAFFT v7.490 (b) Sequences aligned with MUSCLE v5.1. Both trees were inferred using IQ-TREEv1.6.12 with Model Finder limited to LG. Branches are scaled by amino acid substitutions. The red dots indicate the placement of novel tunicate-associated viruses, and support values (sh-aLRT/UFboot) are shown at select nodes.

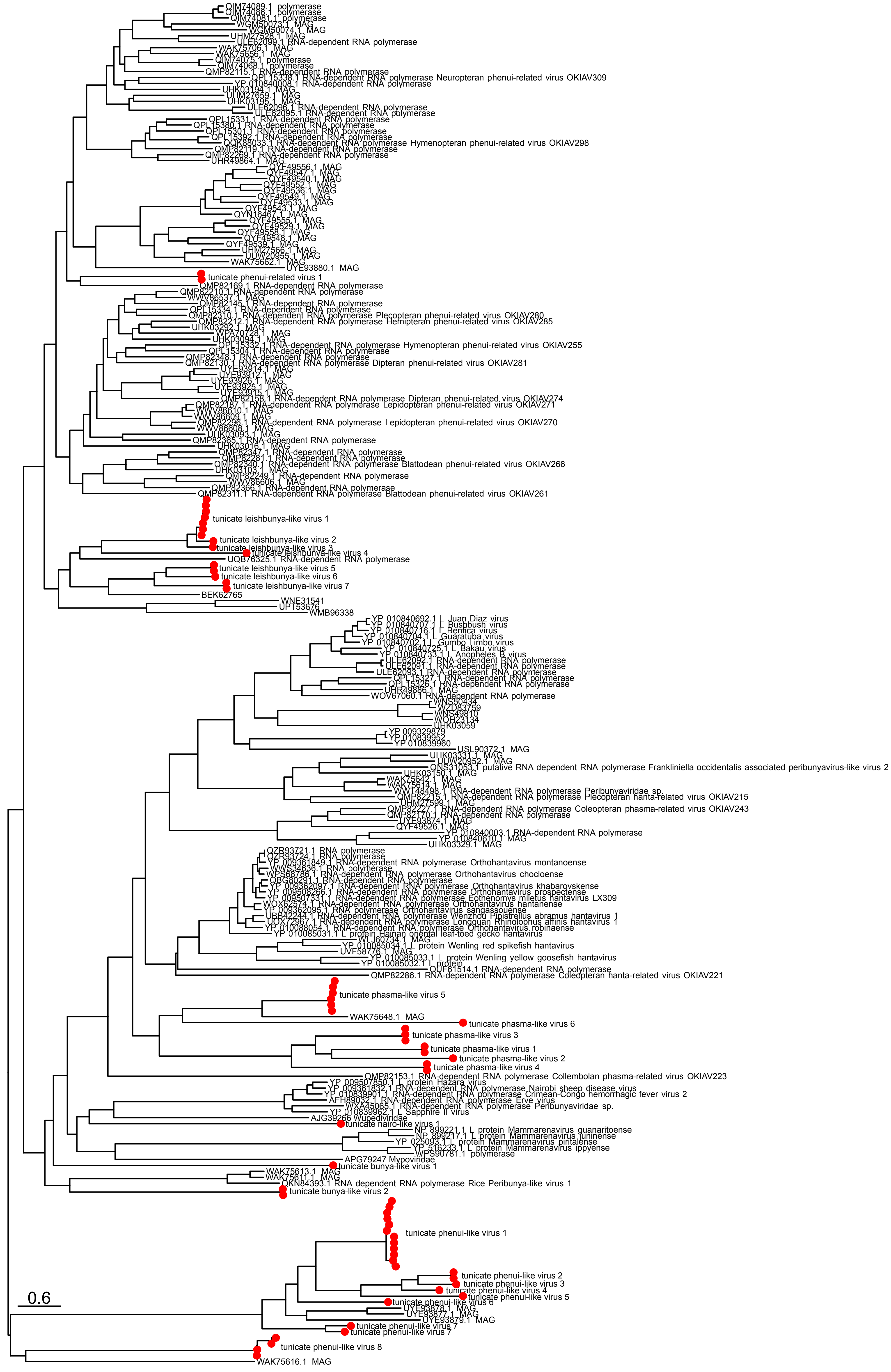

**Figure S4 Midpoint rooted phylogeny of the *Bunyavirales.*** Sequences were aligned with MAFFT v7.490, and the tree was inferred using IQ-TREE v1.6.12 with Model Finder limited to LG. Branches are scaled by amino acid substitutions. The red dots indicate the placement of novel tunicate-associated viruses.

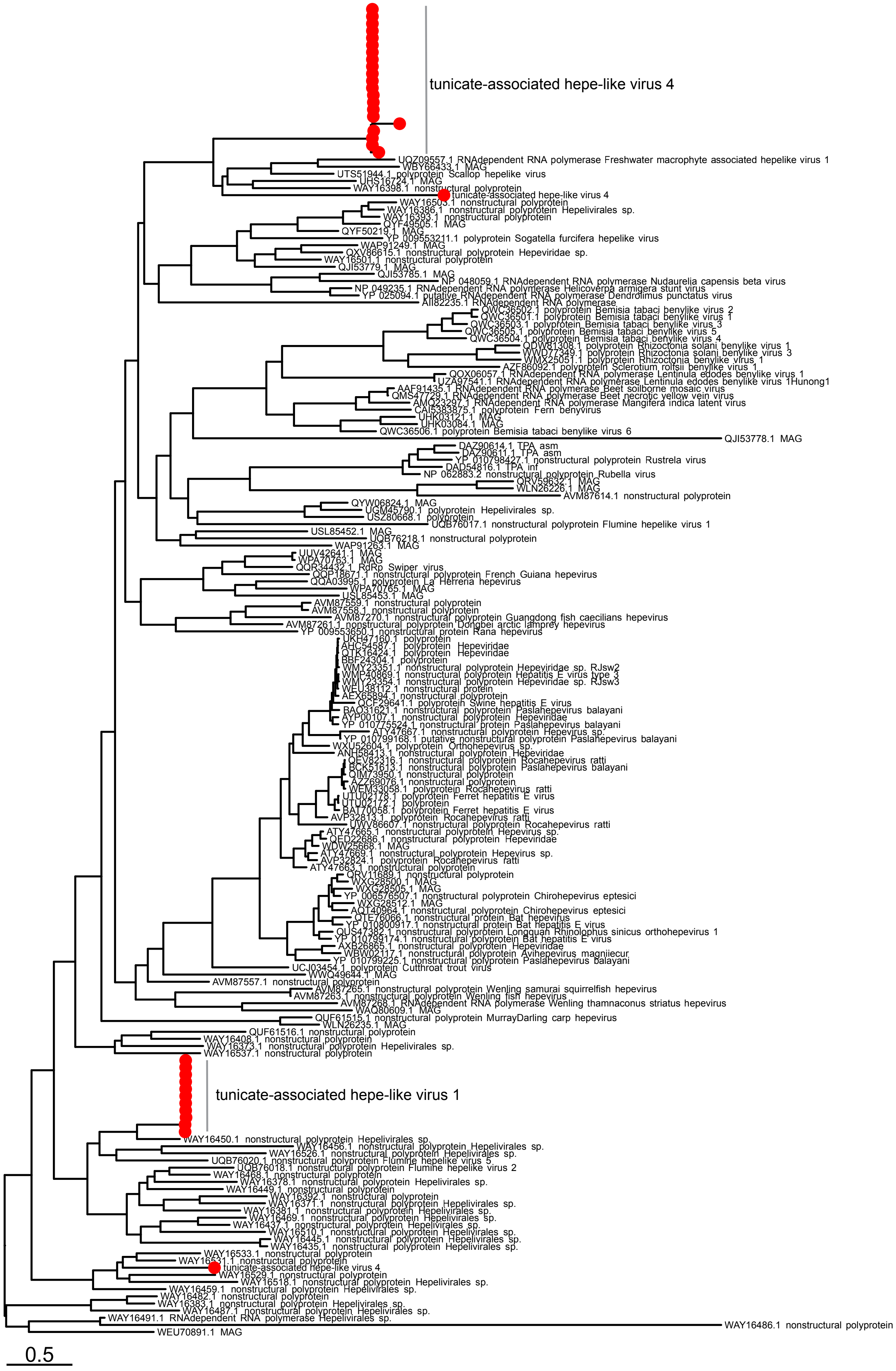


**Figure S5 Midpoint rooted phylogeny of the *Hepelivirales*.** Sequences were aligned with MAFFTv7.490,and the tree was inferred using IQ-TREE v1.6.12 with Model Finder limited to LG. Branches are scaled by amino acid substitutions. The red dots indicate the placement of novel tunicate-associated viruses.

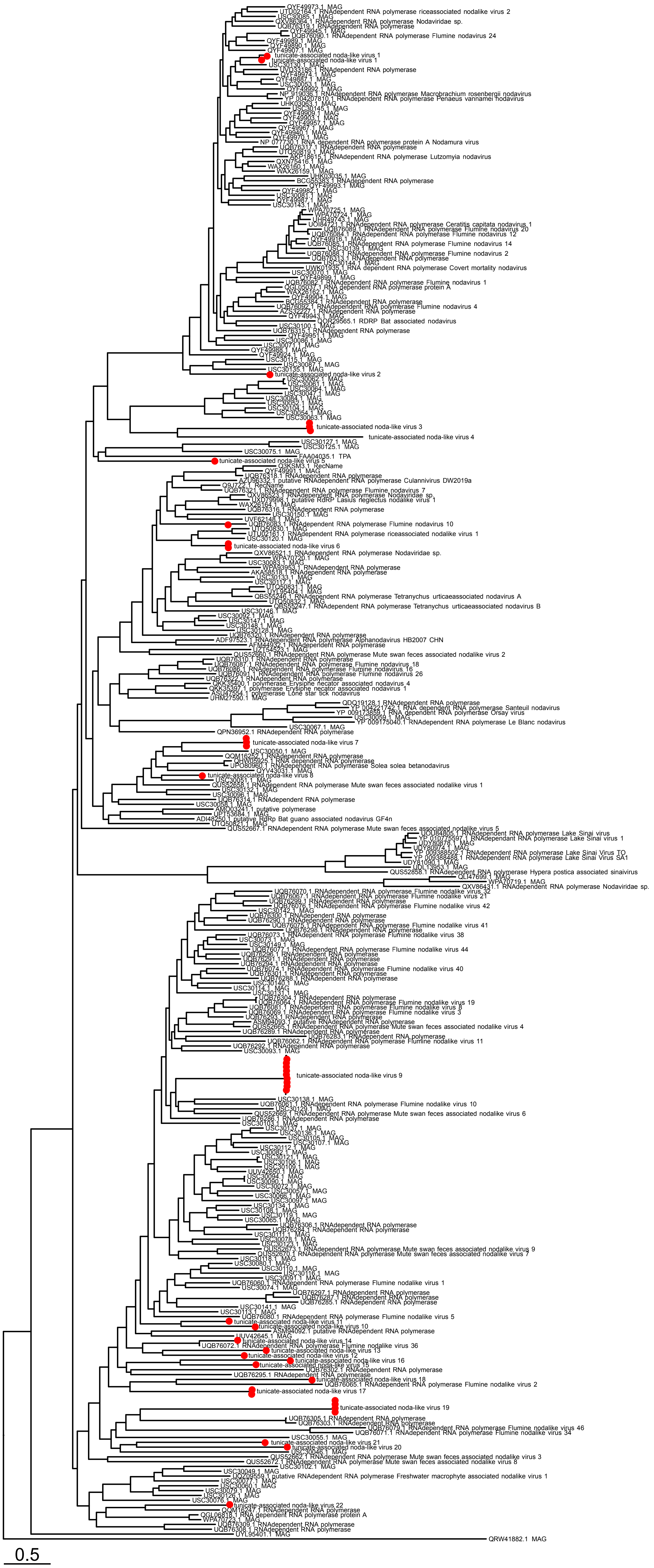


**Figure S6 Midpoint rooted phylogeny of the *Nodamuvirales*.** Sequences were aligned with MAFFT v7.490, and the tree was inferred using IQ-TREE v1.6.12 with Model Finder limited to LG. Branches are scaled by amino acid substitutions. The red dots indicate the placement of novel tunicate-associated viruses.

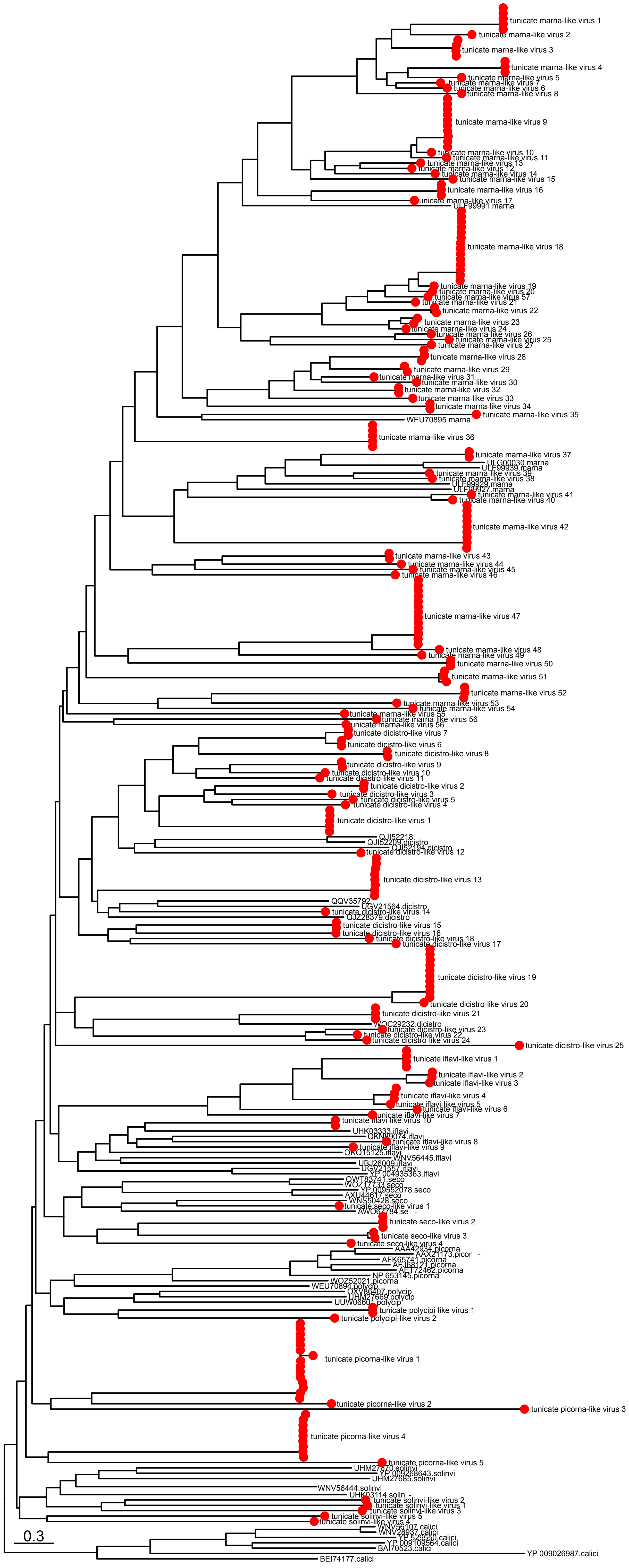

**Figure S7 Midpoint rooted phylogeny of the *Picornavirales.*** Sequences were aligned with MAFFT v7.490,and the tree was inferred using IQ-TREEv1.6.12 with ModelFinder limited to LG. Branches are scaled by amino acid substitutions. The red dots indicate the placement of novel tunicate-associated viruses.

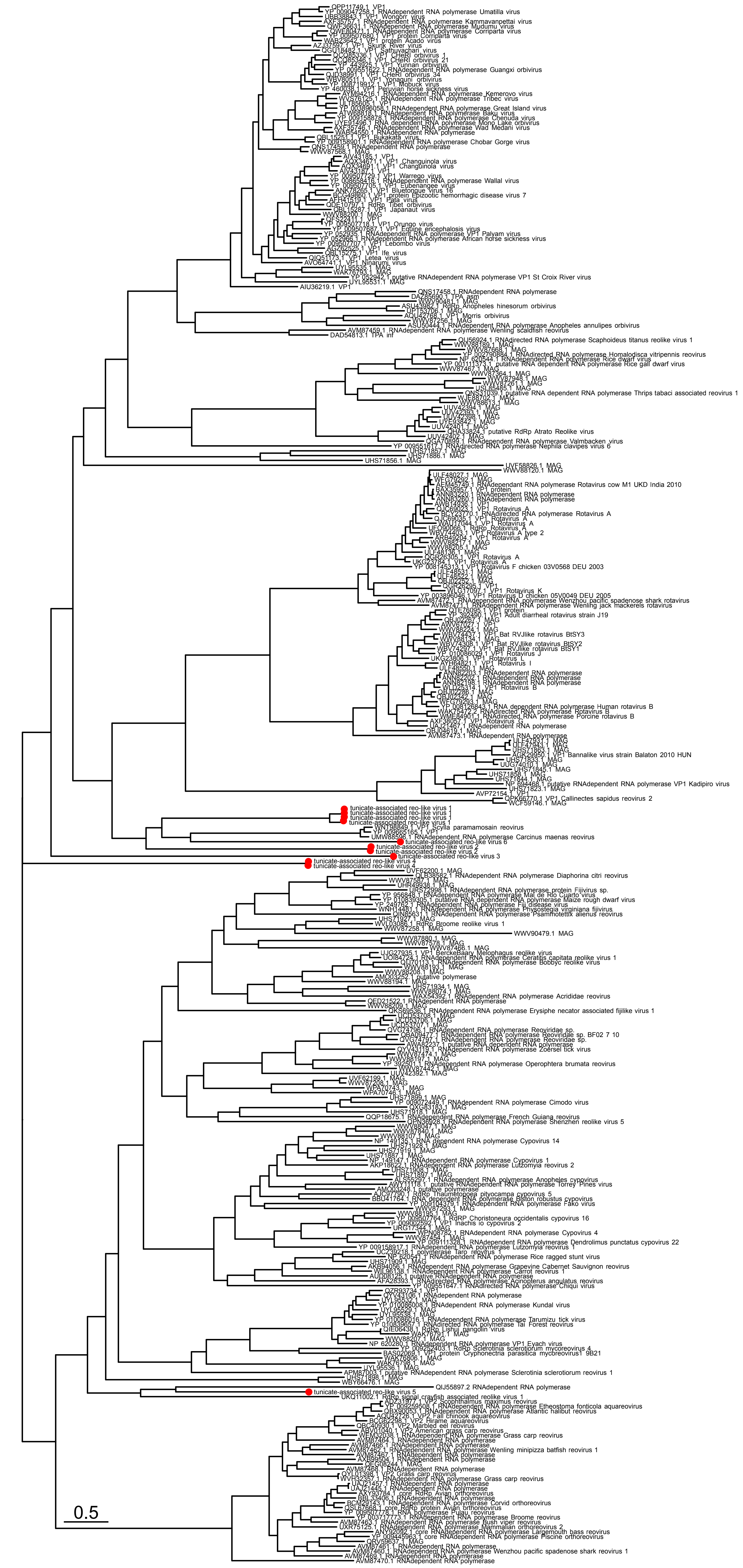

**Figure S8 Midpoint rooted phylogeny of the *Reovirales* polymerase segment.** Sequences were aligned with MAFFT v7.490,and the tree was inferred using IQ-TREE v1.6.12 with Model Finder limited to LG. Branches are scaled by amino acid substitutions. The red dots indicate the placement of novel tunicate-associated viruses.

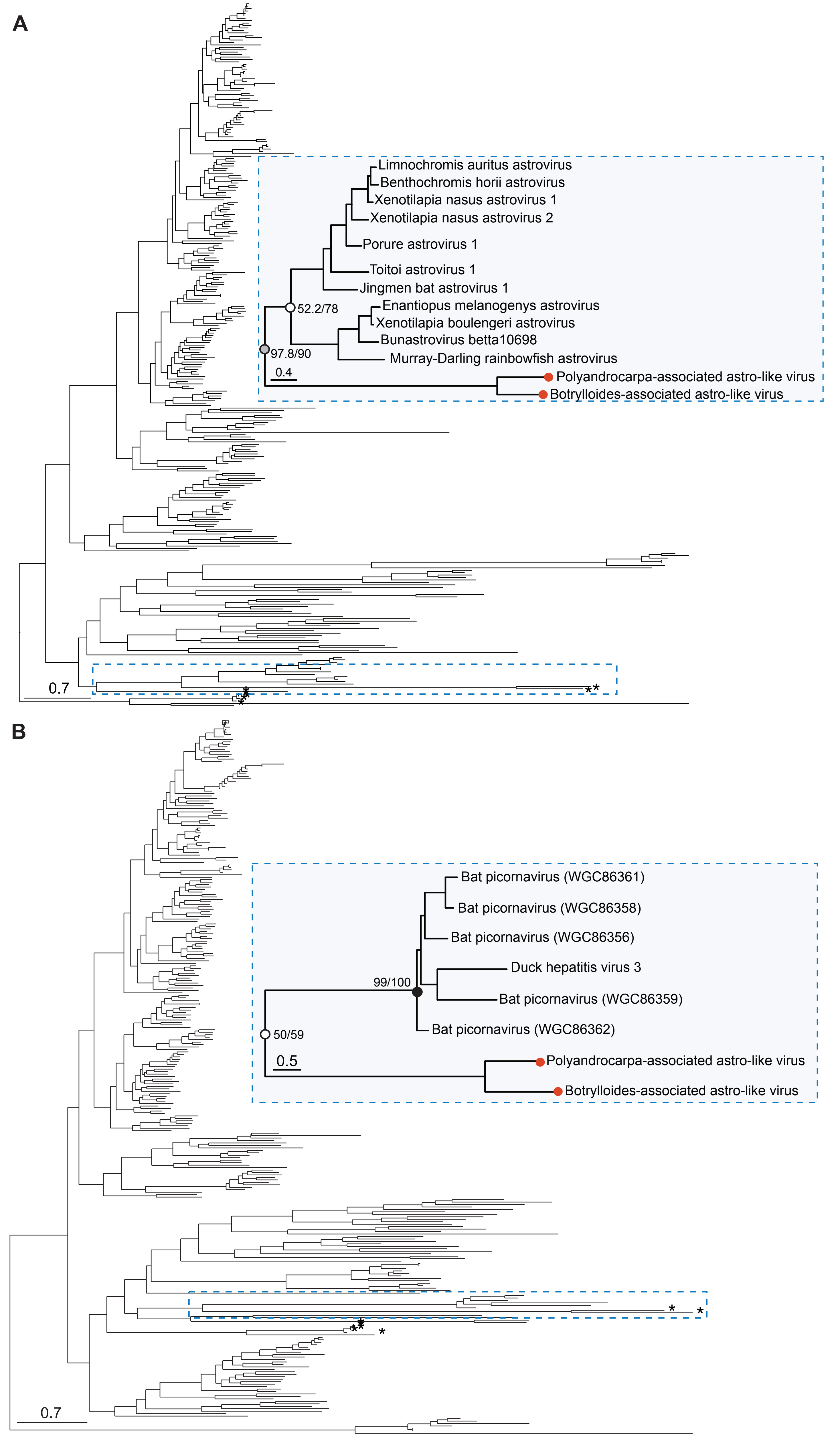


**Figure S9 Midpoint rooted phylogenies of the *Stellavirales*.** (a) Sequences aligned with MAFFTv7.490 (b) Sequences aligned with MUSCLE v5.1. Both trees were inferred using IQ-TREEv1.6.12 with Model Finder limited to LG. Branches are scaled by amino acid substitutions.The stars and red dots indicate the placement of tunicate-associated viruses identified in this study. Support values (sh-aLRT/UFboot) are shown at select nodes.

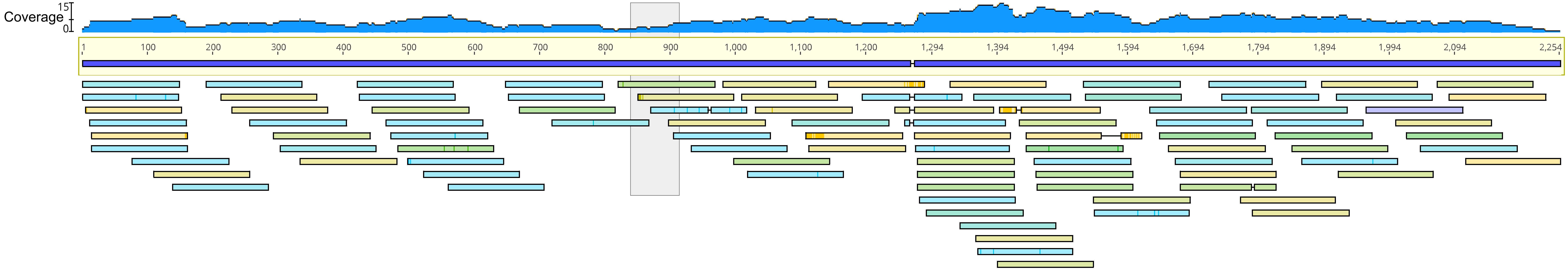


**Figure S10 Sequencing coverage of Clavelina lepadiformis influenza-like virus PB1 segment.** The y-axis shows sequencing coverage, and the grey box indicates the position of the frameshift. The schematic was visualised in Geneious Prime 2025.0.1.

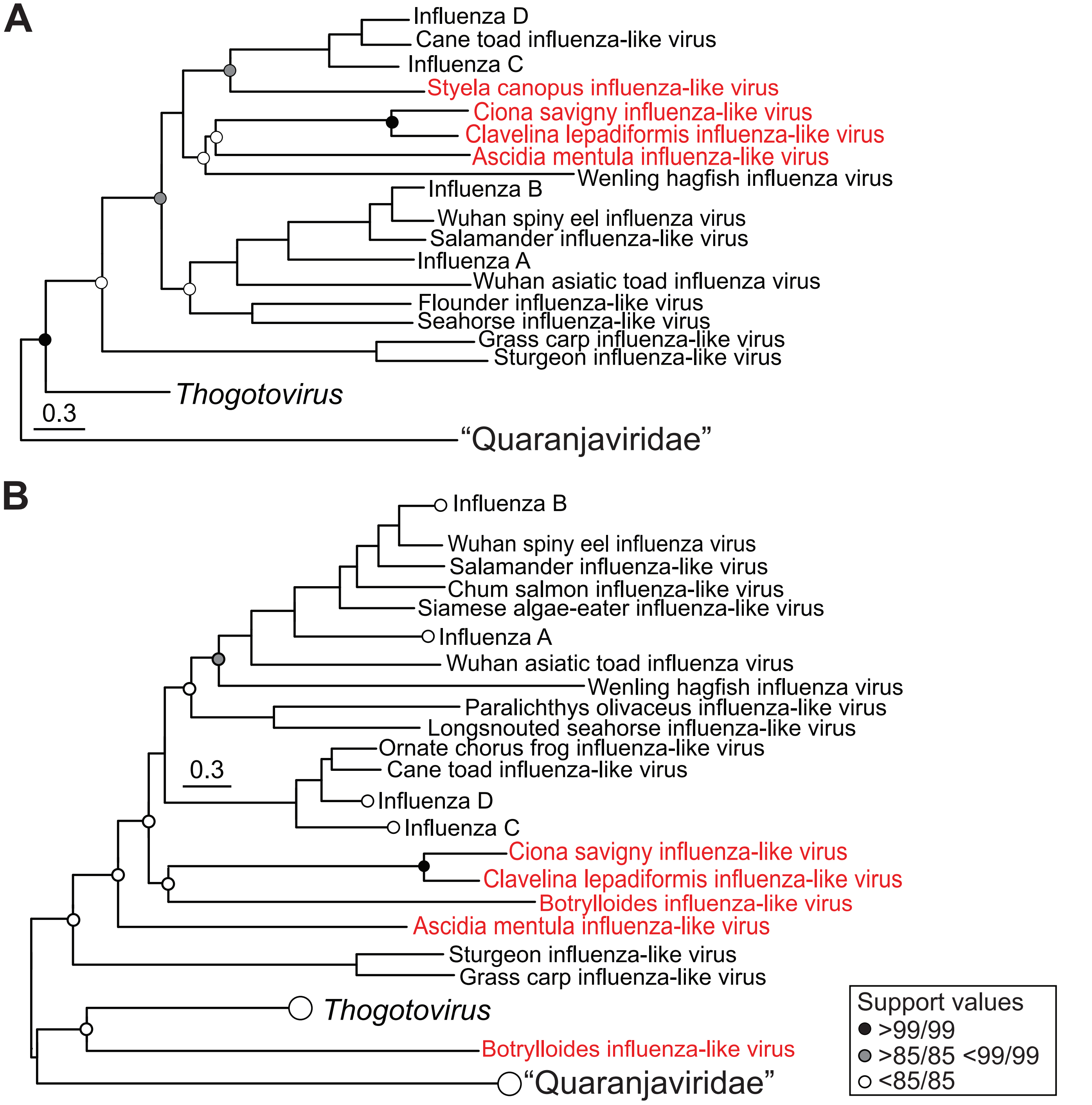


**Figure S11 Phylogenetic inference of influenza PB2 (a) and PB3 (b) segments.** Sequences were aligned with MAFFT and topologies were inferred using IQ-TREE with no restrictions to ModelFinder. Branches are scaled to amino acid substitutions. Red tip labels indicate tunicate-associated segments identified in this study. Support values are shown at select nodes.

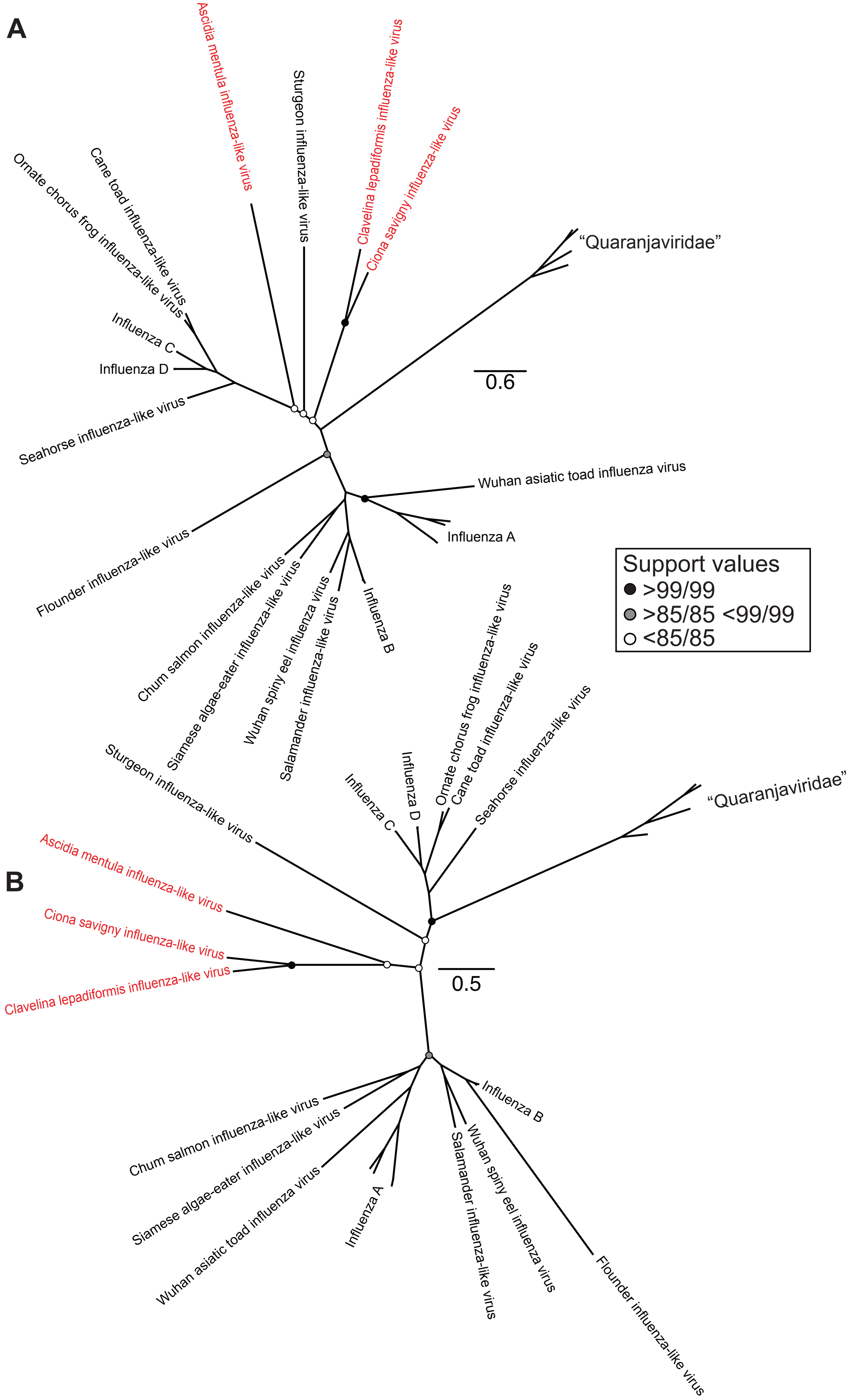


**Figure S12 Unrooted phylogenies of influenza HA segments.** Sequences were aligned with MAFFT (a) or MUSCLE (b). Phylogenies were inferred using IQ-TREE with no restrictions to ModelFinder. Branches are scaled to amino acid substitutions. Red tip labels indicate tunicate-associated segments identified in this study. Support values are shown at select nodes.

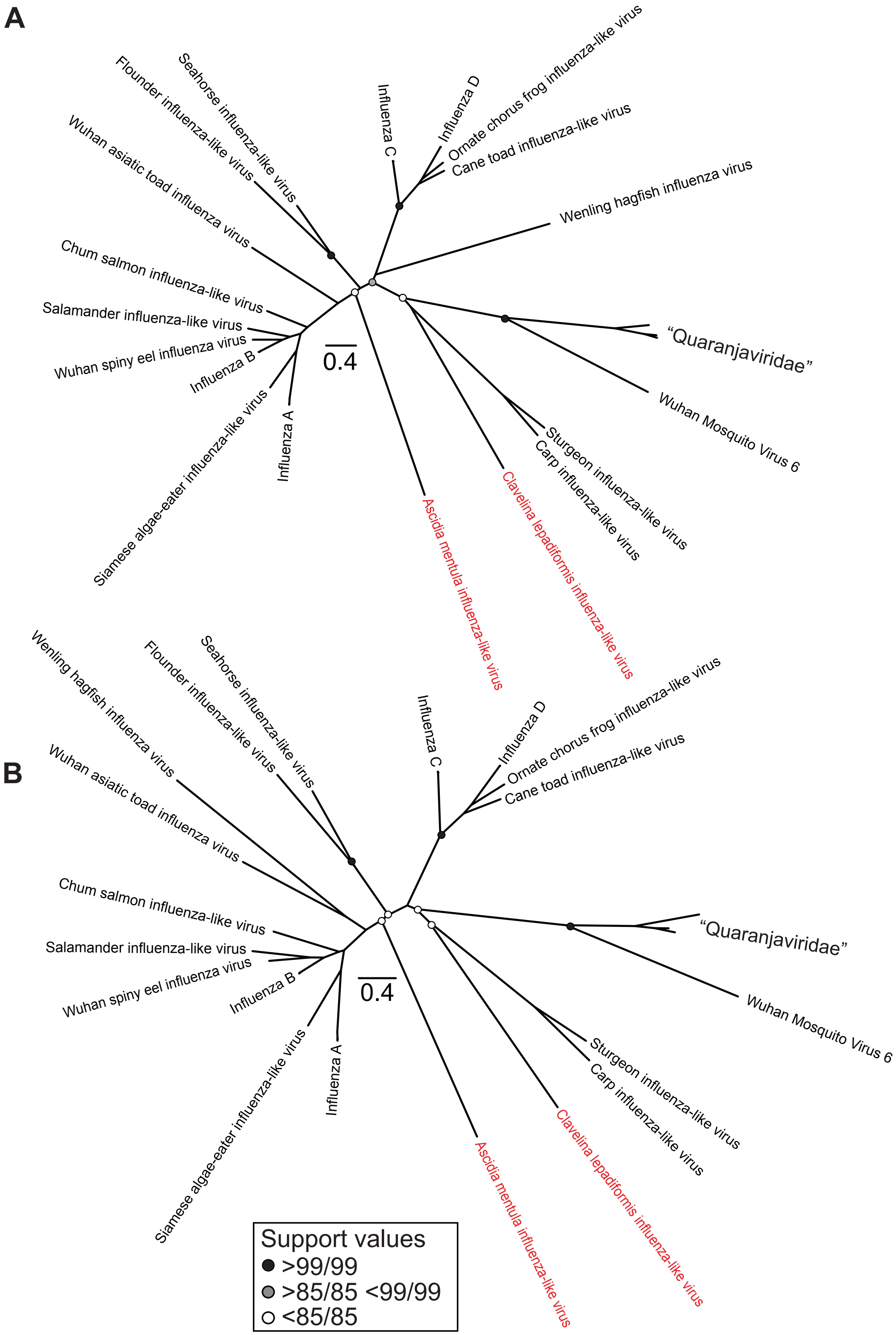


**Figure S13 Unrooted phylogenies of influenza NP segments.** Sequences were aligned with MAFFT (a) or MUSCLE (b). Phylogenies were inferred using IQ-TREE with no restrictions to ModelFinder. Branches are scaled to amino acid substitutions. Red tip labels indicate tunicate-associated segments identified in this study. Support values are shown at select nodes.

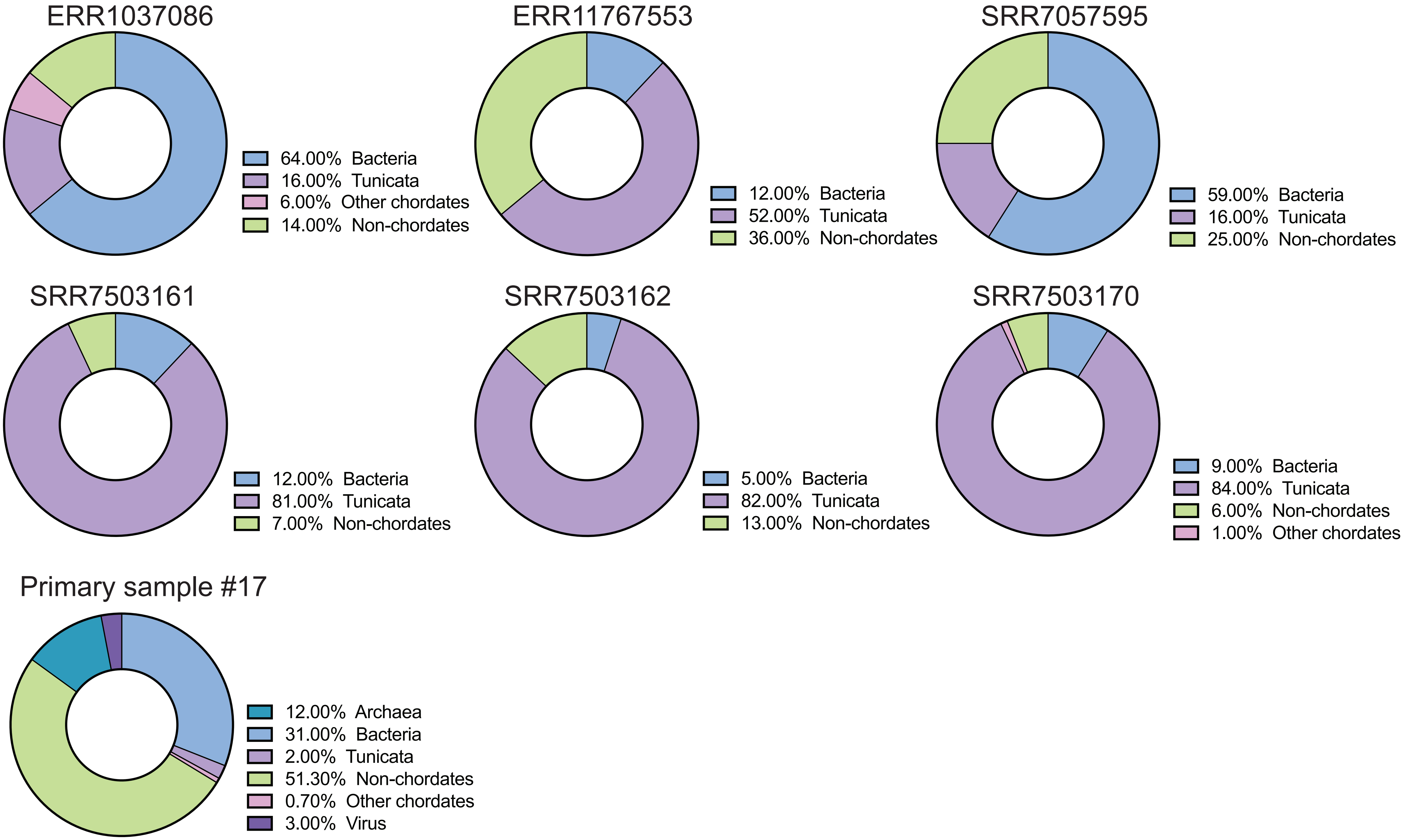


**Figure S14 Library composition of ascidian libraries with influenza-like viral segments.** Libraries were analysed with KMA and CCMetagen. The category "Non-chordates" includes non-tunicate invertebrates and non-animal eukaryotes.

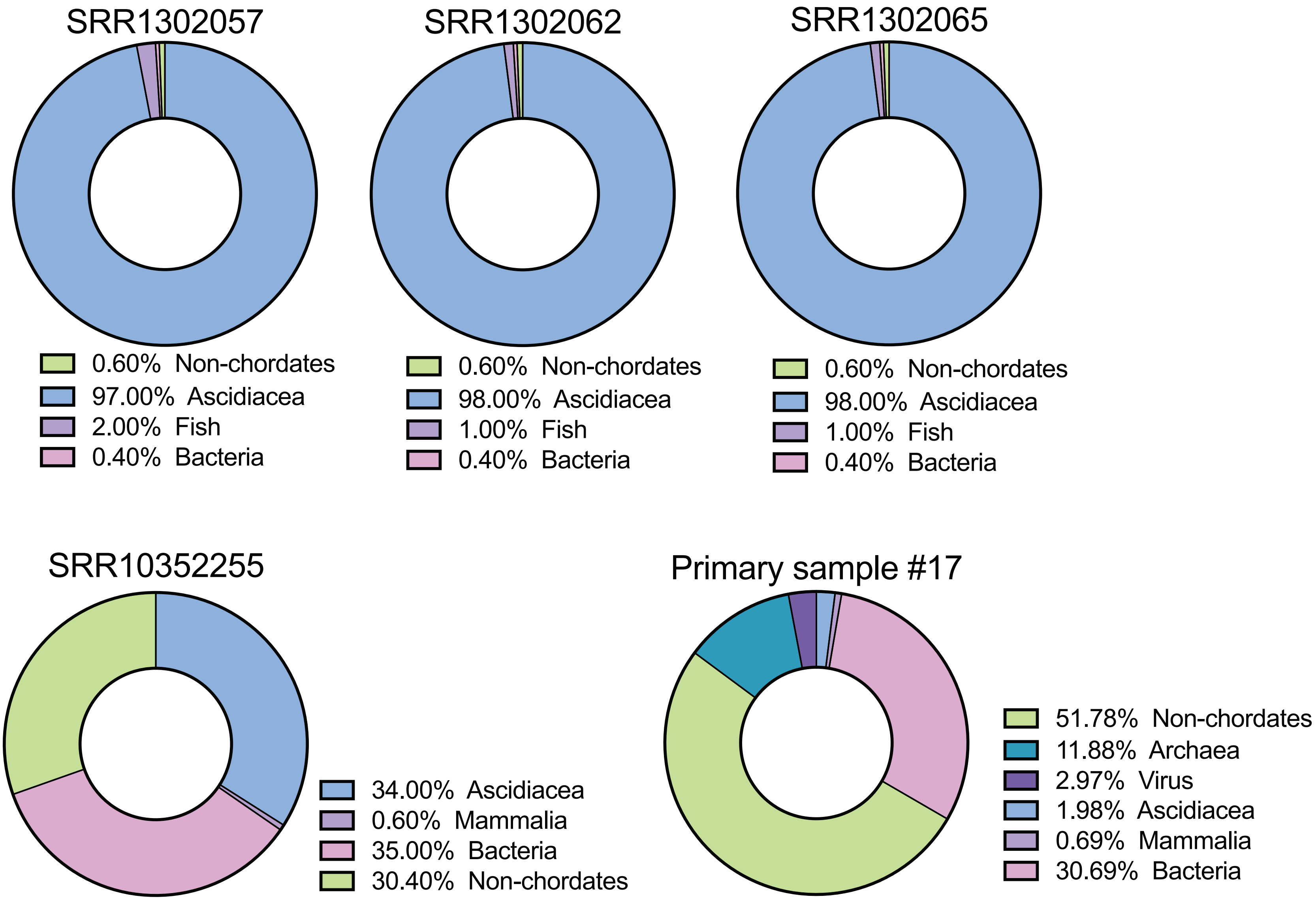


**Figure S15 Library composition of ascidian libraries with novirhabdo-like viruses.** Libraries were analysed with the KMA and CCMetagen. The category “Non-chordates” includes non-tunicate invertebrates and non-animal eukaryotes.

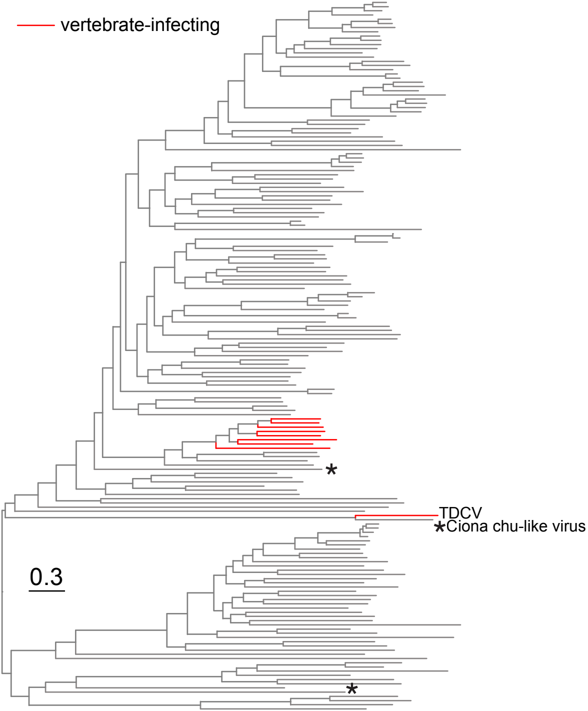


**Figure S16 Midpoint rooted phylogeny of the *Jingchuvirales* RdRp-encoding ORF.** Amino acid sequences were aligned with MUSCLEv5. Branches are scaled to amino acid substitutions. Stars indicate viruses identified in this study. TDCV: Tasmanian devil chu-like virus

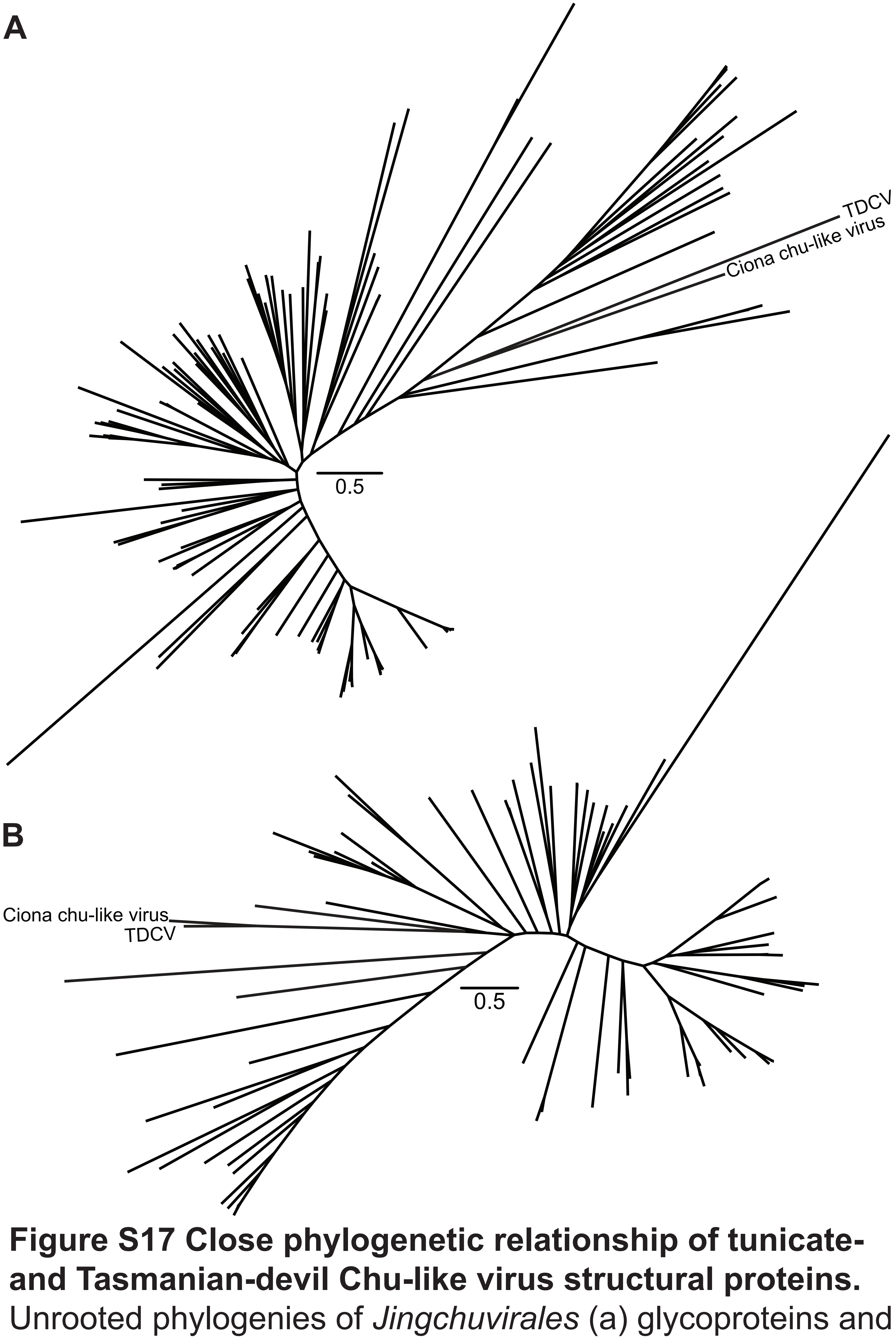


**Figure S17 Close phylogenetic relationship of tunicate- and Tasmanian-devil Chu-like virus structural proteins.** Unrooted phylogenies of *Jingchuvirales* (a) glycoproteins and (b) nucleoproteins. Amino acid sequences were aligned with MAFFT. Branches are scaled by amino acid substitution. TDCV: Tasmanian devil Chu-like virus.

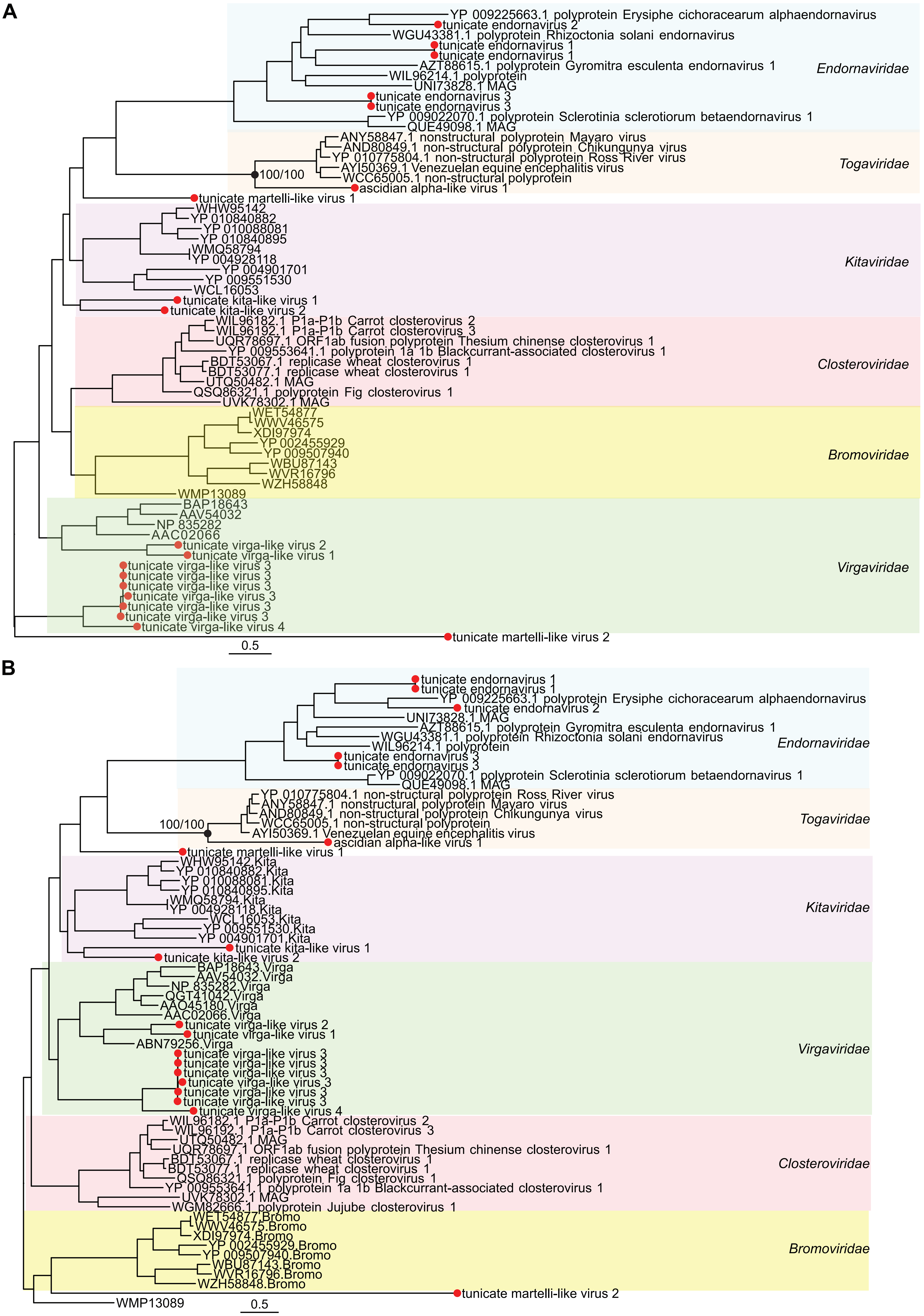


**Figure S18 Midpoint rooted phylogenies of the *Martellivirales* RdRp.** (a) Sequences aligned with MAFFT v7.490 (b) Sequences aligned with MUSCLE v5.1. Both trees were inferred using IQ-TREEv1.6.12 with Model Finder limited to LG. Branches are scaled by amino acid substitutions.The red dots indicate the placement of tunicate-associated viruses identified in this study, and support values (sh-aLRT/UFboot) are shown at select nodes.

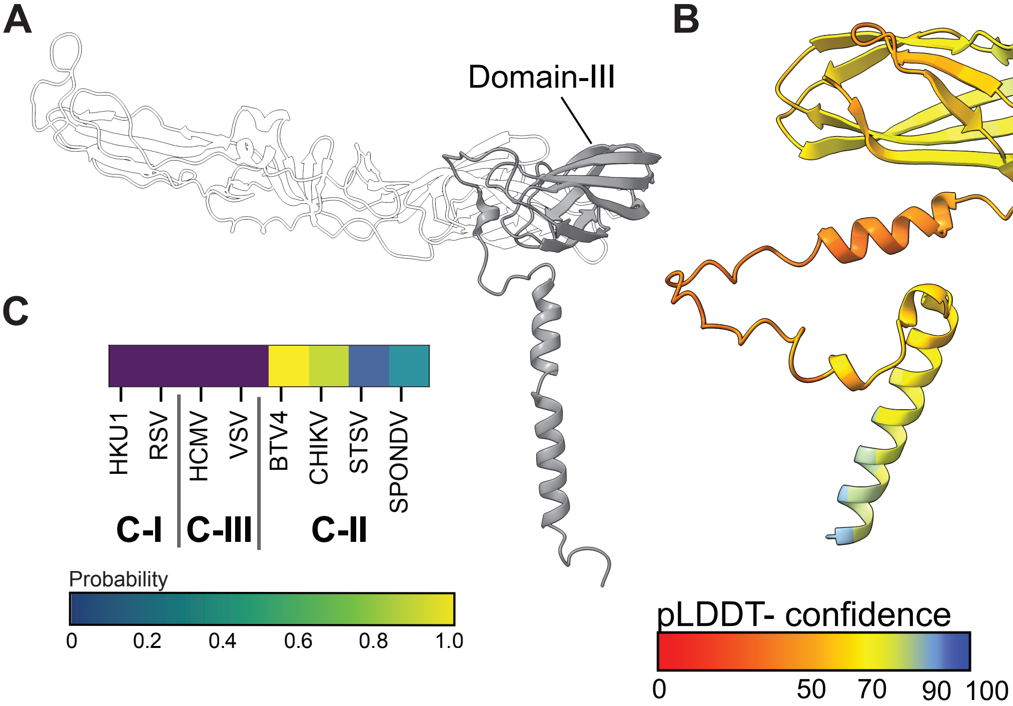


**Figure S19 Flatworm monono-like virus glycoprotein is Class II-like.**
(a) Structure of Chikungunya E1 Glycoprotein (Reference PDB 6NK7). Domain-III is highlighted in dark grey. (b) ESMFold predicted structure of Domain-III of Flatworm monono-like virus glycoprotein. Protein structures for sequential overlapping sequence blocks, spanning entire viral polyproteins, were compared to class-I (C-I), III (C-III) and II (C-II) fusion protein reference structures using Foldseek. The ribbon diagrams is color-coded by prediction confidence (pLDDT) (c) Heat map displays lowest detected Foldseek e-values against each fusion protein reference.

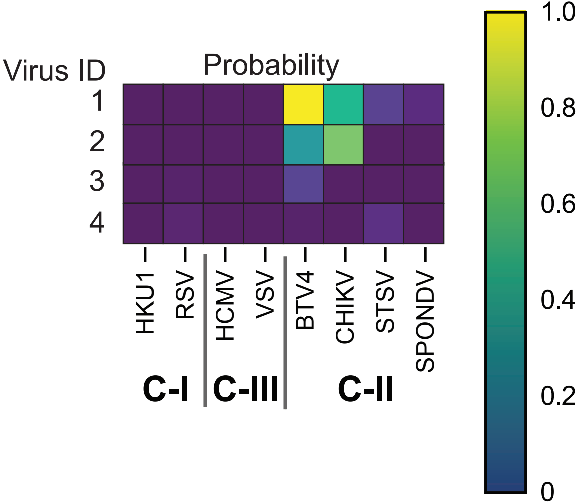


**Figure S20. FoldSeek probabilities of structural homology of divergent nido-like viruses.** Number key: 1 Supergroup012--nido_soil_SRR5829926_k141_103166_flag1_multi14_len47250, 2 Supergroup012--AI-nido_marine_ERR1719354_k119_30219_flag1_multi570_len19927, 3 Supergroup012--nido_wastewater_SRR8840828_k141_118584_flag1_multi25_len36514, 4 Supergroup012--nido_soil_SRR5215306_k141_264356_flag1_multi30_len38030. Abbreviations: C-I Class I, C-II Class 2, C-III Class 3

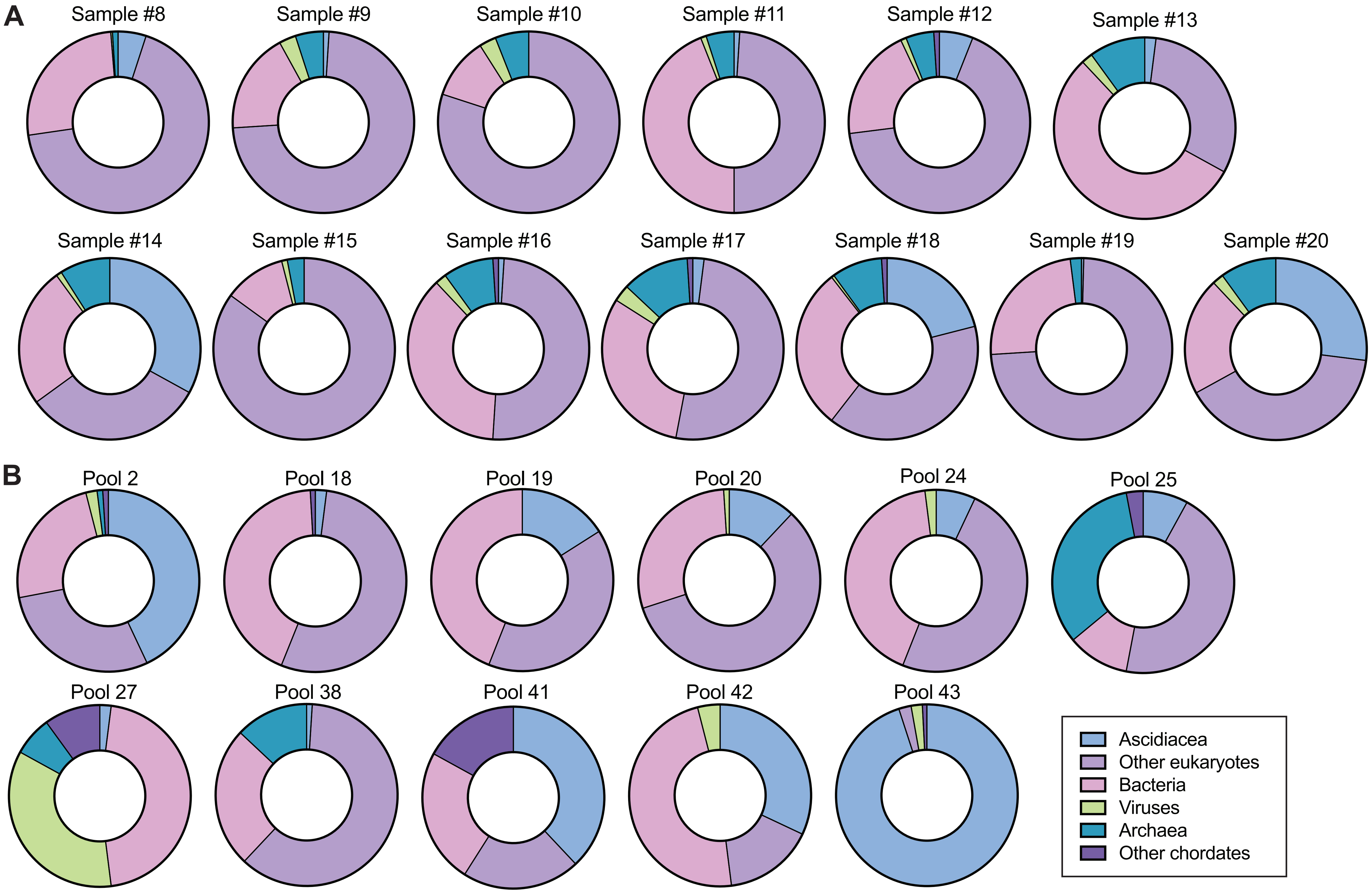


**Figure S21. Library composition of primary libraries.** Samples collected at timepoint 1 (a) and 2 (b).
