## Supplementary Tables S1-S3 for "Tunicate metatranscriptomes reveal ancient virus-host co-divergence and inter-order recombination in the evolutionary history of disease-causing viruses"

**Table S1. HHPred results for ronivirus and euronivirus glycoproteins**

| Input (Family) | HHPred reference name | E-value | Probability (%) |
|---|---|---|---|
| Yellow head virus (*Roniviridae*) | SFTSV, virion | 1.6 | 97.03 |
| Gill-associated virus (*Roniviridae*) | SFTSV, virion | 1.5 | 97.06 |
| Macrobrachium rosenbergii Golda virus (*Roniviridae*) | La Crosse fusion protein | 2.7e-76 | 100 |
| Beihai Nido-like virus 2 (*Euroniviridae*) | SFTSV, virion | 18 | 88.76 |
| Wenling Nido-like virus 1 (*Euroniviridae*) | SFTSV, virion | 6.5 | 90.82 |

SFTSV: Severe Fever with Thrombocytopenia Syndrome Virus (*Bunyavirales*)

**Table S2 HHPred results for glycoproteins encoded by divergent nidoviruses identified using artificial intelligence[36]**

| Glycoprotein ID | HHPred reference name | E-value | Probability (%) |
|---|---|---|---|
| Supergroup012--AI-nido_marine_ERR1719354_k119_30219_flag1_multi570_len19927_ORF2 | Envelopment polyprotein; SFTSV, virion | 130 | 70.44 |
| Supergroup012--nido_soil_SRR5215306_k141_264356_flag1_multi30_len38030_ORF4 | Glycoprotein C; Crimean-Congo hemorrhagic fever virus | 9.2 | 83.36 |
| Supergroup012--nido_soil_SRR5829926_k141_103166_flag1_multi14_len47250_ORF3 | Schmallenberg Virus Envelope Glycoprotein | 120 | 63.88 |
| Supergroup012--nido_wastewater_SRR8840828_k141_118584_flag1_multi25_len36514_ORF2 | Glycoprotein C; Crimean-Congo hemorrhagic fever virus | 0.14 | 96.78 |

### Table S3 Length thresholds used to screen virus candidates

| Order | Length cut-off (nt) |
|---|---|
| *Amarillovirales* | 1000 |
| *Articulavirales* | 500 |
| *Birnaviridae* | 500 |
| *Bunyavirales* | 1000* |
| *Hepelivirales* | 1000 |
| *Martellivirales* | 1000 |
| *Mononegavirales* | 3000† |
| *Nidovirales* | 4000*‡ |
| *Nodamuvirales* | 1000 |
| *Picornavirales* | 3000 |
| *Reovirales* | 500 |
| *Stellavirales* | 1000 |

*When the RdRp catalytic triad (e.g., SDD) was included
†Except when there were multiple, smaller fragments in a single library
‡For SRA libraries; 1000nt was used for primary libraries
